# Robust Cancer Mutation Detection with Deep Learning Models Derived from Tumor-Normal Sequencing Data

**DOI:** 10.1101/667261

**Authors:** Sayed Mohammad Ebrahim Sahraeian, Li Tai Fang, Marghoob Mohiyuddin, Huixiao Hong, Wenming Xiao

## Abstract

Accurate detection of somatic mutations is challenging but critical to the understanding of cancer formation, progression, and treatment. We recently proposed NeuSomatic, the first deep convolutional neural network based somatic mutation detection approach and demonstrated performance advantages on *in silico* data. In this study, we used the first comprehensive and well-characterized somatic reference samples from the SEQC-II consortium to investigate best practices for utilizing deep learning framework in cancer mutation detection. Using the high-confidence somatic mutations established for these reference samples by the consortium, we identified strategies for building robust models on multiple datasets derived from samples representing real scenarios. The proposed strategies achieved high robustness across multiple sequencing technologies such as WGS, WES, AmpliSeq target sequencing for fresh and FFPE DNA input, varying tumor/normal purities, and different coverages (ranging from 10× - 2000×). NeuSomatic significantly outperformed conventional detection approaches in general, as well as in challenging situations such as low coverage, low mutation frequency, DNA damage, and difficult genomic regions.

## Introduction

Somatic mutations are critical cancer drivers. Accurate somatic mutation detection enables precise diagnosis, prognosis, and treatment of cancer patients^1^. Several tools have been developed to identify somatic mutations from next-generation sequencing technology^2–11^. In general, conventional techniques pinpoint somatic mutations from background noise, germline variants, and/or cross-contaminations through various statistical/algorithmic modeling approaches employing a set of hand-crafted sequencing features extracted in the tumor-normal paired sequencing data^2–10^. Those manually designed approaches are not easily generalizable beyond the specific cancer types, sample types, or sequencing strategies for which they were developed.

Recently, a deep learning based somatic mutation detection approach, called NeuSomatic, has been proposed that uses a convolutional neural network (CNN) to learn feature representations directly from raw data^11^. NeuSomatic uses a novel summarization of tumor/normal alignment information as a set of input matrices that can be adopted to efficiently train models that learn how to effectively differentiate true somatic mutations from artifacts. The network models trained by NeuSomatic can capture important mutation signals directly from read alignment and genomic-context without manual intervention. NeuSomatic, thus, provides a framework that can easily be applied to different problem statements including diverse sequencing technologies, cancer types, tumor and normal purities, and mutation allele frequencies, through training on proper data. Moreover, it can be implemented as a stand-alone somatic mutation detection method or with an ensemble of existing methods to achieve the highest accuracy. NeuSomatic has been shown to significantly outperform conventional techniques on *in silico* data sets. Given the lack of thoroughly-characterized benchmark samples with known “ground-truth” somatic mutations, performance evaluation on real samples was limited to a partial sensitivity analysis on a small number of validated somatic variants. Thus, despite the advantages seen by implementing NeuSomatic’s CNN-based framework on *in silico* data, the accuracy and reproducibility of NeuSomatic on real cancer samples have not been comprehensively evaluated to date.

Recently, the Somatic Mutation Working Group in the FDA-led Sequencing Quality Control Phase II (SEQC-II) consortium developed reference matched tumor-normal samples: a human triple-negative breast cancer cell line (HCC1395) and a matched B lymphocyte derived normal cell line (HCC1395BL)^12,13^. Using orthogonal sequencing technologies, multiple sequencing replicates, and multiple bioinformatics analysis pipelines, the SEQC-II consortium has developed a well-defined “Gold Set” of somatic Single Nucleotide Variants (SNVs) and insertion/deletions (INDELs) for HCC1395.

As the first comprehensive and well-characterized paired tumor-normal reference cancer samples, this dataset along with the accompanying sequencing data prepared on multiple sites and technologies provides a unique resource to achieve two important purposes. First, using the high-confidence somatic mutation call set developed for these reference samples by the consortium, we performed in-depth analysis of deep learning based somatic mutation detection in a real cancer sample, in comparison with the conventional schemes. Second, we explored various model-building strategies using SEQC-II data to train the CNN in NeuSomatic and identified effective training approaches on multiple data sets derived from samples representing real scenarios. We evaluated the proposed strategies on whole genome sequencing (WGS), whole exome sequencing (WES), and AmpliSeq targeted sequencing datasets with converges ranging from 10 × − 2000 ×. The WGS and WES data were derived from Formalin-Fixed Paraffin-Embedded (FFPE) and fresh DNA, using three library preparation protocols with various input amounts, and from multiple platforms/sequencing sites. Using tumor and normal titration, we evaluated different approaches for 5%- 100% tumor purities, 5% of contaminated match normal, and with 10 × - 300 × WGS coverage. The proposed strategies to train and implement the deep learning framework in NeuSomatic achieved high robustness across all the aforementioned real scenarios, and significantly outperformed conventional paired tumor-normal somatic mutation detection approaches.

Our analysis on SEQC-II reference cancer samples demonstrates that the deep learning scheme implemented in NeuSomatic helps to overcome the main challenges in somatic mutation detection which are not easy to resolve using conventional techniques. The deep learning models and strategies derived from our study can thus provide the research community with actionable best-practice recommendations for robust cancer mutation detection.

## Results

### Reference samples and datasets

For full-spectrum analysis of the somatic mutation detection problem, we used the first comprehensive whole-genome characterized reference tumor-normal paired breast cancer cell lines (HCC1395 and HCC1395BL), developed by the Somatic Mutation Working Group of the SEQC-II consortium^12,13^. We leveraged high-confidence somatic mutations (39,536 SNVs and 2,020 INDELs) derived by the consortium as our ground truth set (**Suppl. Figure 1**). For a broad assessment of consistency and reproducibility of predictions, we used a total of 123 replicates from diverse datasets which represent realistic cancer detection applications including real WGS, WES, and AmpliSeq target sequencing with different coverages, tumor purities, and library preparations, on FFPE and fresh DNA sequenced with multiple platforms in six centers (See Methods).

### Analysis overview

We evaluated NeuSomatic’s deep learning based somatic mutation detection approach using various trained network models and compared it with seven widely-used conventional somatic mutation calling algorithms including MuTect2^2^, MuSE^3^, SomaticSniper^4^, Strelka2^5^, VarDict^6^, TNscope^7^, and Lancet^8^. We assessed NeuSomatic both in its standalone mode (shown as NeuSomatic-S) and its ensemble mode, where predictions reported by MuTect2, SomaticSniper, VarDict, MuSE, and Strelka2 are also included as input channels in addition to raw data.

We used several different training models in our analysis. First, we used the already available model published recently^11^ which was trained using *in silico* spike-ins from the DREAM Challenge Stage 3 dataset^13^. Despite the large discrepancy between the sample types, sequencing platforms, coverages, spike-in mutation frequencies, and heterogeneity of the samples used to train the DREAM3 model, this model outperformed other conventional techniques across the real cancer datasets of diverse characteristics by more than ∼4% by the mean F1-score averaged across different samples for both SNVs and INDELs (**Figure 1a**). Although this superiority supports the robustness of NeuSomatic to the stated variations, it also suggests that the deep learning framework used in NeuSomatic could perform even better through learning the sequencing features and mutation signatures from real cancer samples, especially for predicting INDELs, and for higher coverage PCR-enriched datasets like AmpliSeq.

**Figure 1.**
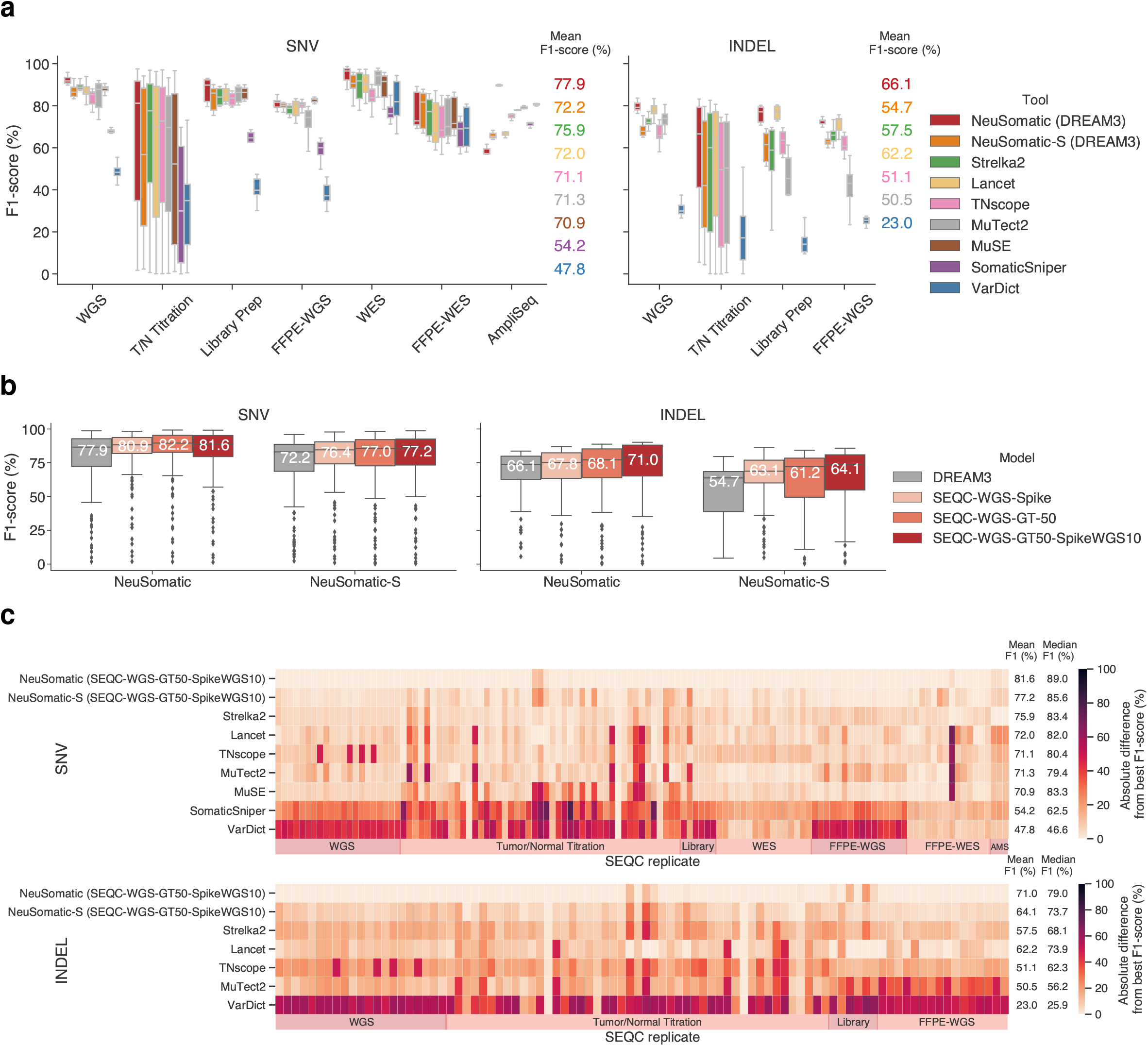
NeuSomatic’s overall performance on 123 replicates in SEQC-II dataset. (a) The NeuSomatic model trained on DREAM challenge stage 3 dataset outperformed other techniques when applied to SEQC-II data. (b) Training different NeuSomatic models using SEQC-II reference samples resulted in additional absolute 4-5% improvement in the mean F1-score. (c) Using model trained on SEQC-II data, NeuSomatic achieves consistent superiority over other techniques across diverse set of replicates of different purities/coverages in WGS, WES, FFPE, AmpliSeq, and different library-prep datasets. In this subfigure, for each replicate the best F1-score was computed across different approaches. The heatmaps illustrate the absolute difference between the F1-score of any of the somatic mutation detection approaches to the best F1-score. In each panel, the mean F1-score is shown for each approach across 123 replicates.

To identify the most effective strategies for building network models using real cancer samples, we evaluated ten more training approaches for NeuSomatic using the SEQC-II reference samples (**Suppl. Table 1**). The first model (SEQC-WGS-Spike) was trained on a set of WGS pairs of *in silico* tumor-normal replicates, where the *in silico* tumors were designed by spiking mutations into distinct normal replicates. The second model (SEQC-WGS-GT-50) was trained using the ground truth somatic mutations in HCC1395 on a set of real WGS tumor-normal replicate pairs on 50% of the genome. The third model (SEQC-WGS-GT50-SpikeWGS10) was prepared by adding 10% of the training data from the first model to those for the second model to take advantage of both a high number of spike-in mutations and the realistic somatic mutations. These three models were tested on all datasets. For the specific datasets like FFPE and WES, we also prepared six additional specialized models using a set of synthetic pairs of *in silico* tumor and normal replicates. For all models, we evaluated the performance on the 50% held-out region of the genome (not used for the SEQC-WGS-GT-50 model). We also trained a model (SEQC-WGS-GT-ALL) using all ground truth mutations similar to the SEQC-WGS-GT-50 model but on the whole genome. Rather than being directly applicable to our WGS dataset where it has seen all true mutations, SEQC-WGS-GT-ALL will be beneficial for performance analysis on other datasets or samples other than HCC1395.

Employing the models trained on SEQC-II samples, we boosted the average DREAM3 model performance by an additional mean F1-score improvement of ∼4-5% (**Figure 1b**). The proposed model-building strategy was consistently best across various sample types and sequencing strategies, outperforming conventional approaches by more than 5.7% and 7.8% in terms of the mean F1-score for SNVs and INDELs, respectively. Similarly, we observed more than 5.6% improvement compared to the median F1-score of other conventional techniques across all samples (**Figure 1c**; **Suppl. Figure 2**).

### WGS dataset

We evaluated the performance of the aforementioned somatic calling techniques and network models on the 21 WGS replicates sequenced in six sequencing centers using the HiSeqX10, HiSeq4000, and NovaSeq S6000 platforms (**Figure 2; Suppl. Table 2**). The NeuSomatic SEQC-WGS-GT50-SpikeWGS10 model performed consistently better than other schemes with minor variations across replicates, demonstrating robustness and reproducibility (**Figure 2a**). NeuSomatic yielded average F1-scores of 94.6% and 87.9% for SNVs and INDELs with more than 5.6% and 10.2% superiority, respectively in SNVs and INDELs over the mean F1-scores of other conventional somatic mutation detection schemes. Precision-recall analysis revealed that the high precision of NeuSomatic compared to other techniques drove this superiority (**Figure 2b**). Comparing different model training strategies also revealed that NeuSomatic-S INDEL calling benefit more from training using ground truth somatic mutations, while, in general, we observed significant improvements of up to 11% from using SEQC-II reference samples compared to the DREAM3 model (**Figure 2c**).

**Figure 2.**
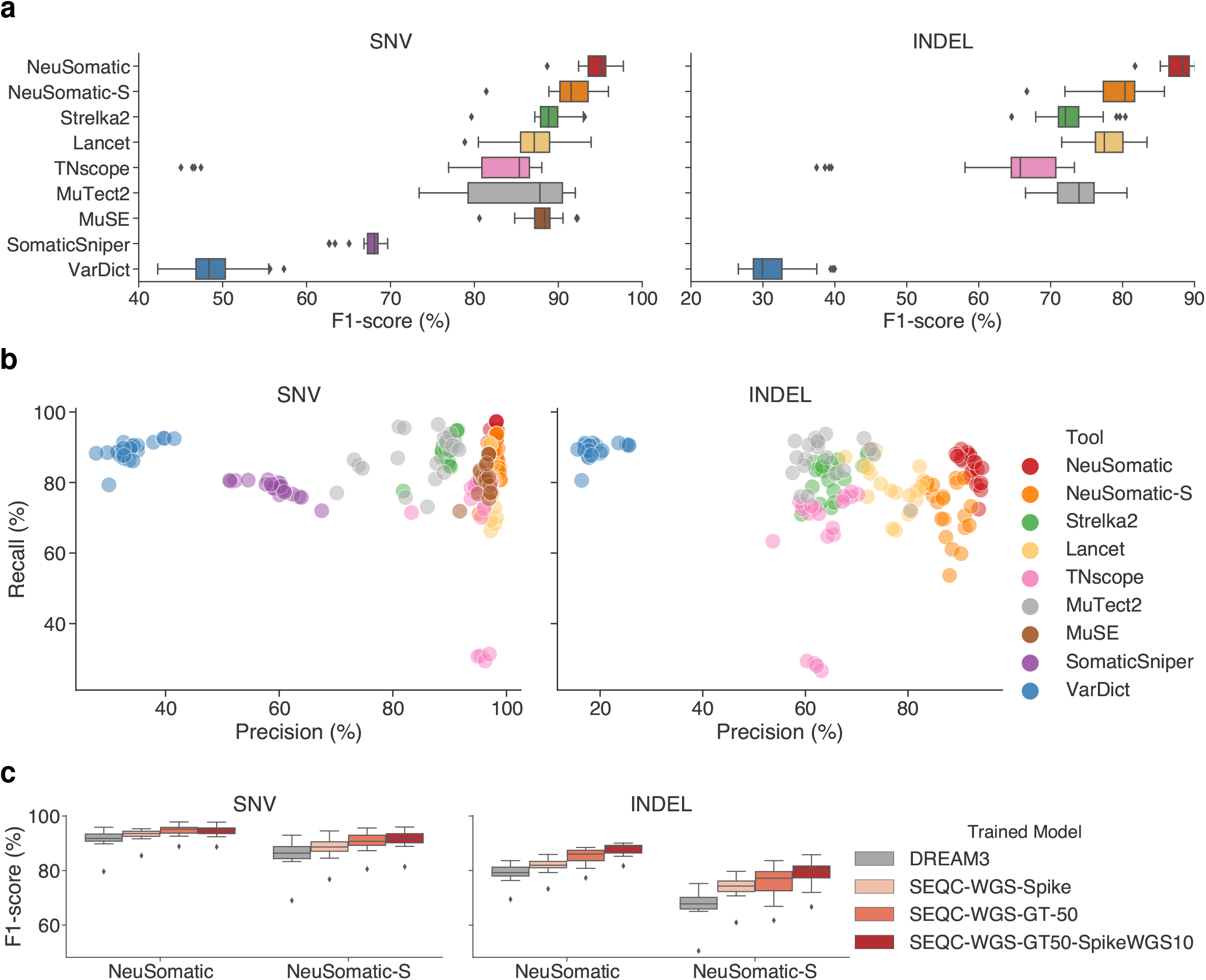
Performance comparison on 21 WGS replicates dataset. (a) F1-score (%) and (b) Precision-Recall comparison across different somatic mutation callers. Here, the SEQC-WGS-GT50-SpikeWGS10 trained models were used for NeuSomatic and NeuSomatic-S. (c) F1-score (%) comparison for various NeuSomatic trained models.

### Tumor purity and contaminated normal

As tumor purity and the contamination of tumor cells in match normal sample would greatly impact mutation detection accuracy, we investigated model robustness on different sequencing depths and sample purities by tumor-normal titration of WGS samples. We first studied tumor samples with 5%-100% purity paired with pure normal samples for 10×-300× coverage ranges (**Figure 3; Suppl. Figure 3 and 4; Suppl. Table 3**). In general, NeuSomatic maintained superiority over other schemes despite tumor purity and coverage variations, which reflected its robustness (**Figure 3a; Suppl. Figure 3**). It had the largest advantages over conventional schemes for more challenging cases like lower coverage (e.g. ∼20% F1-score advantage for a sample with 10× coverage and 100% purity).

**Figure 3.**
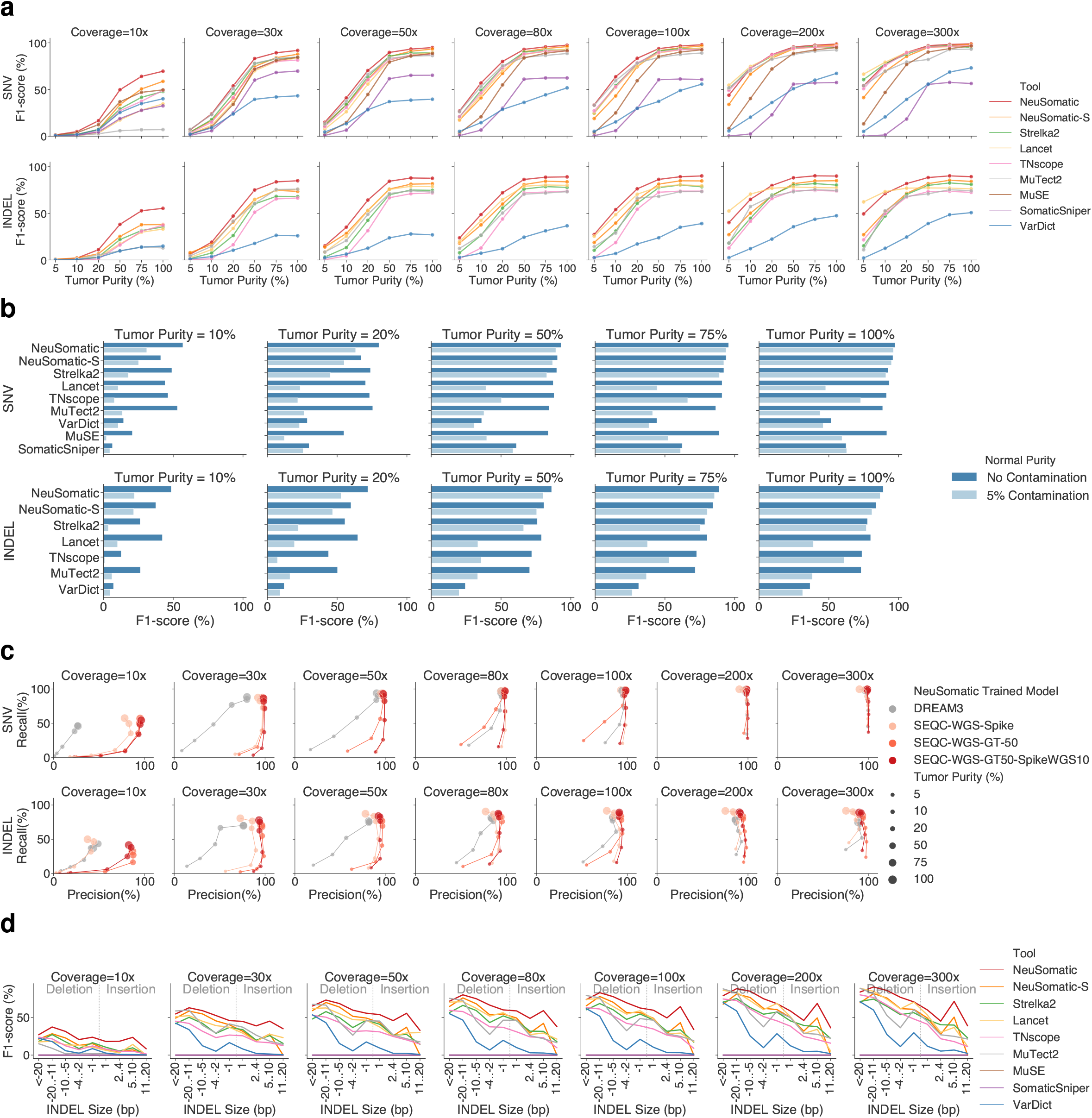
Performance comparison on Tumor-purity dataset. (a) SNV and INDEL F1-score (%) comparison for different somatic mutation callers across different coverages (10×-300×) and tumor purities (5%-100%). (b) Robustness to tumor contamination in the matched normal: F1-score (%) change when matched normal was contaminated with 5% of tumor is shown versus the F1-score (%) at pure normal for 80× coverage and 5-100% tumor purities. For (a) and (b) panels, the SEQC-WGS-GT50-SpikeWGS10 trained models were used for NeuSomatic and NeuSomatic-S. (c) F1-score (%) comparison for various NeuSomatic trained models. (d) Performance analysis for different INDEL sizes. Negative INDEL sizes reflect deletion.

We further analyzed the robustness to 5% tumor contamination in the normal sample for tumor sample purities ranging from 10%-100% at 80x coverage (**Figure 3b**). NeuSomatic yielded high robustness to tumor-normal cross-contamination with less than 5% median absolute change in F1-score. Among the other techniques with high F1-score at pure normal, Strelka2, also showed high robustness to tumor contamination (8.4% median change in F1-score). MuTect2, MuSE, Lancet, and TNscope, despite having high F1-scores for the pure normal scenario, experienced significant drops of up to ∼50% in F1-score when contaminated normal was used.

The models trained on the ground truth variants yielded the highest advantage over the DREAM3 model, in general, mainly due to the higher precision (**Figure 3c; Suppl. Figure 4**). The SEQC-WGS-Spike model, trained on *in silico* tumors, also showed lower precision compared to models trained on the ground truth. In general, INDELs, low purity, and low coverage samples benefited the most from training on SEQC-II data.

Analyzing the F1-scores for different INDEL sizes across various coverages and purity settings revealed NeuSomatic’s robustness to size variations for both insertions and deletions (**Figure 3d**).

### Library-preparation and DNA input

To measure the impact of library preparation on prediction robustness, we used our models to test the six replicates prepared using TruSeq-Nano and Nextera Flex protocols and three DNA input amounts: 1ng, 10ng, and 100ng (**Figure 4; Suppl. Table 4**). NeuSomatic consistently outperformed other techniques across different library preparation approaches. For the 1 ng TruSeq-Nano libraries, all the methods had poor performance due to the limited effective coverage after removal of redundant reads (∼7x). On average, NeuSomatic yielded 8.4% F1-score improvement for SNVs over conventional somatic mutation detection techniques. For INDELs, Lancet’s assembly-based INDEL caller outperformed the NeuSomatic SEQC-WGS-GT50-SpikeWGS10 model by ∼4% (**Figure 4a**). In contrast, NeuSomatic’s SEQC-WGS-GT50 model achieved similar performance to Lancet for INDELs (**Figure 4b, Suppl. Figure 5**). Training on SEQC-II spike-in or ground truth data resulted in overall ∼8.4% F1-score improvement for SNVs compared to DREAM3 model. The advantage was more pronounced in more challenging cases including TruSeq-Nano libraries with 1ng input. We also observed that NeuSomatic-S benefited more from these models.

**Figure 4.**
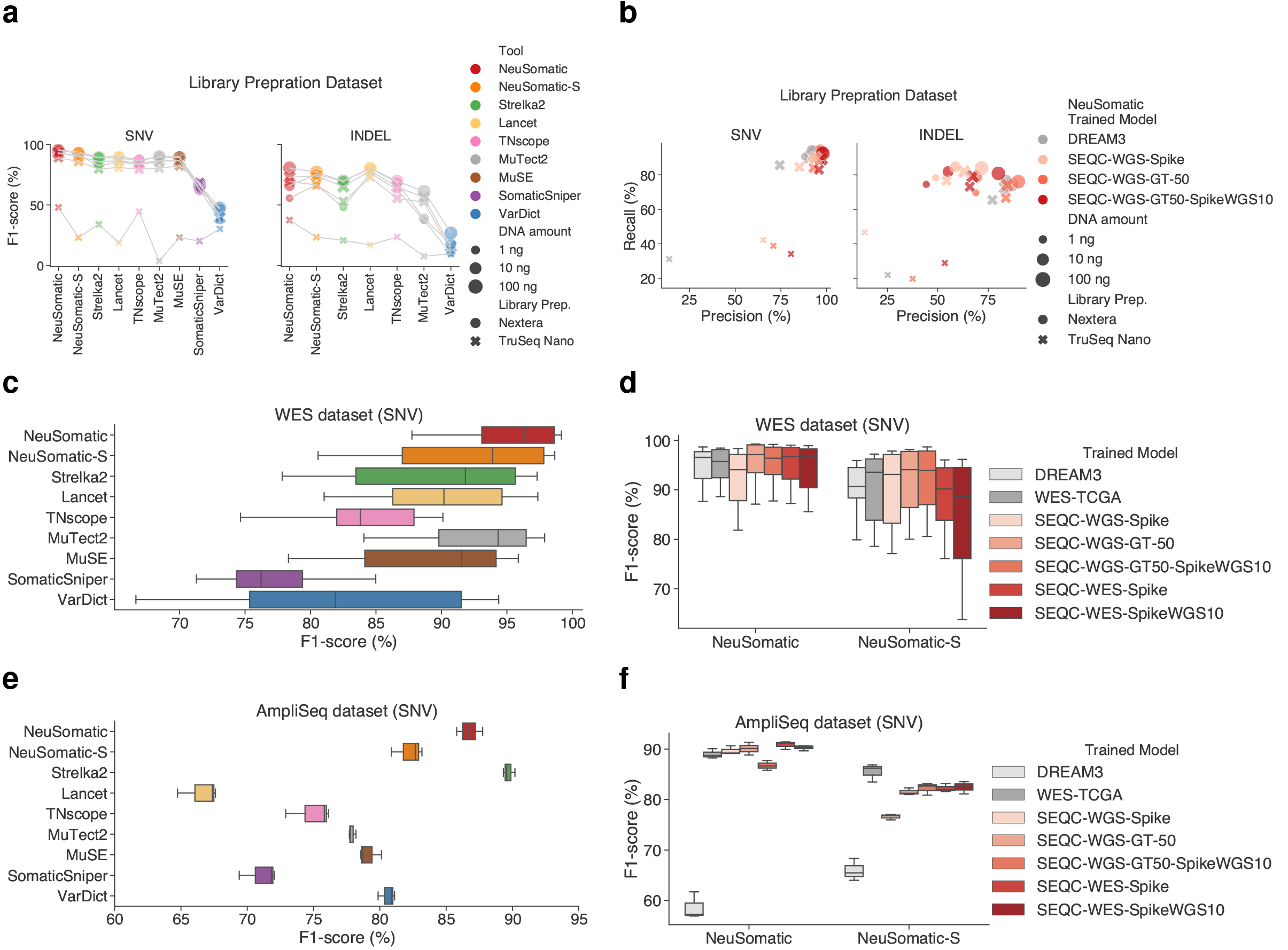
Performance comparison of library prep, WES, and AmpliSeq datasets. (a) SNV and INDEL F1-score (%) comparison for different somatic mutation callers across different library-kits and DNA amounts. (b) Precision-Recall comparison for various NeuSomatic trained models for different library-kits and DNA amounts. (c, e) SNV F1-score (%) comparison for different somatic mutation callers. (d, f) F1-score (%) comparison for various NeuSomatic trained models. For (a), (c), and (e) panels, the SEQC-WGS-GT50-SpikeWGS10 trained models were used for NeuSomatic and NeuSomatic-S.

### Captured (WES) and Targeted (AmpliSeq) panels

We tested our models on the 16 WES replicates sequenced at six sequencing sites, as well as three replicates in AmpliSeq dataset (**Figures 4c-f; Suppl. Tables 5 and 6).** Although the models trained on SEQC-II WGS samples had different coverage and platform biases, NeuSomatic performed well on both WES and AmpliSeq datasets with 2000x coverage. For WES, NeuSomatic achieved an average SNV F1-score of 95.4% with more than 2.6% improvement over the mean F1-score of alternative schemes. On the WES dataset, models trained on WES and WGS data performed similarly with ∼95% F1-score. On the AmpliSeq dataset, the NeuSomatic SEQC-WG-GT50 and SEQC-WES-Spike models achieved average F1-scores of greater than 90%, which were the highest compared to other models/schemes along with Strelka2.

### Effect of FFPE processing

To measure the robustness of NeuSomatic’s prediction on FFPE processed samples, we used 8 WGS and 7 WES FFPE replicates, prepared with four different formaldehyde fixation times of 1 hr, 2 hrs, 6 hrs, and 24 hrs. We evaluated each FFPE replicate with both FFPE and fresh normal matched sample. NeuSomatic continued consistently superior performance over the other techniques despite presence of FFPE artifacts and remained largely invariant to fixing time and the matched normal sample used (**Figure 5a; Suppl. Tables 7 and 8**). On WGS FFPE data, NeuSomatic yielded average F1-scores of 86.1% and 76.9%, respectively for SNVs and INDELs, with more than 4% and 6% improvement over the mean F1-score of alternative techniques. Similarly, for FFPE WES data, NeuSomatic yielded an average F1-score of 78.9%, with more than 4% superiority over the mean F1-score of conventional schemes (**Figure 5c**).

**Figure 5.**
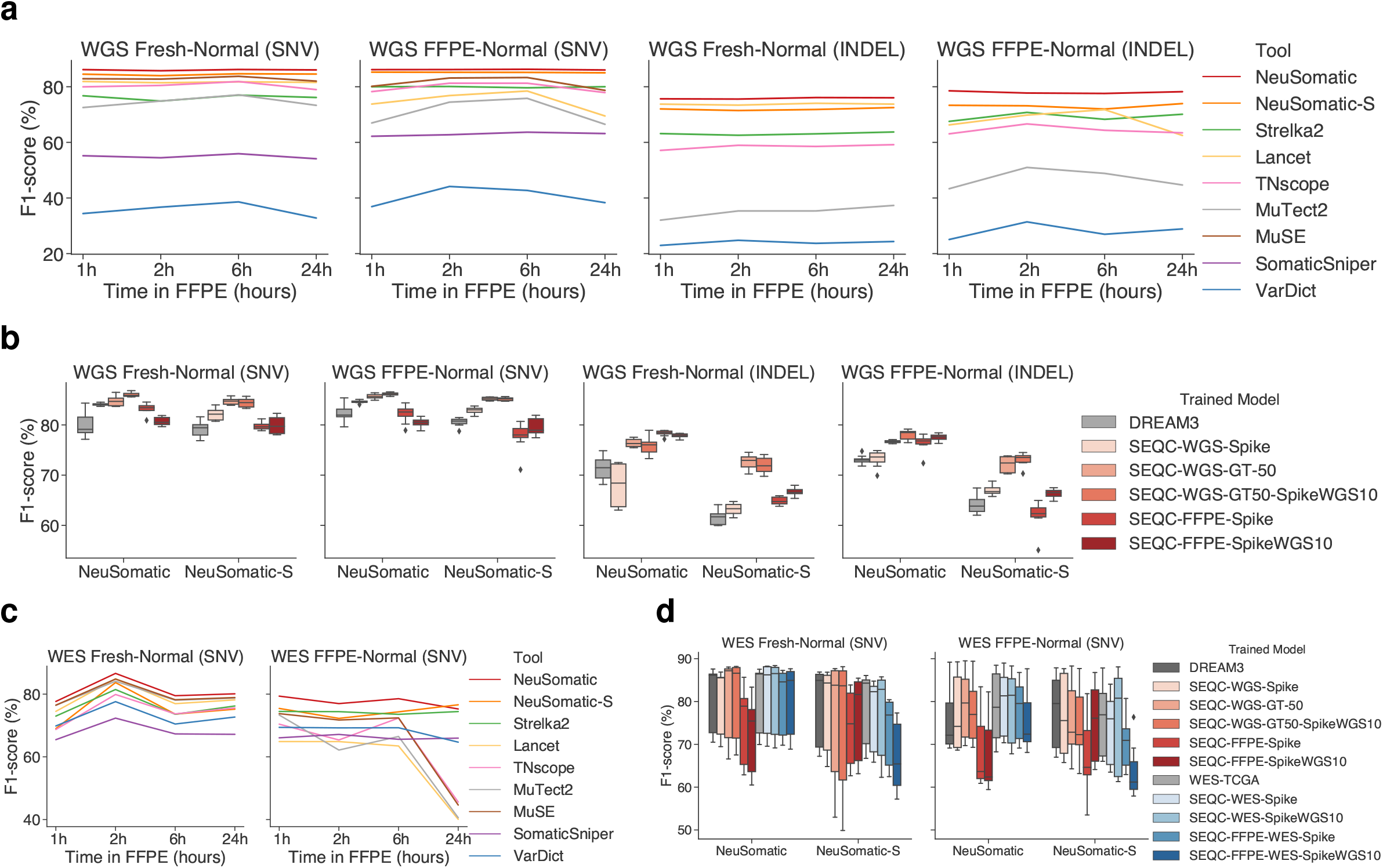
Performance comparison of FFPE dataset. (a, b, c) Performance on 16 FFPE WGS replicates. (d, e) Performance on 14 FFPE WES replicates. (a, d) F1-score (%) comparison for different somatic mutation callers across FFPE and fresh normal samples. (b, e) F1-score (%) comparison for various NeuSomatic trained models. (c) Precision-Recall plot of 16 FFPE WGS replicates for different somatic mutation callers with FFPE and fresh normal. For (a) and (c) panels, the SEQC-WGS-GT50-SpikeWGS10 trained models were used for NeuSomatic and NeuSomatic-S.

In general, we observed significant boost over the DREAM3 model when we leveraged SEQC-II samples for training (**Figures 5b and 5d**). The models trained on FFPE samples seemed to improve those trained on fresh sample, only for INDEL prediction using NeuSomatic, but, for SNVs, models trained on fresh samples were superior.

### Sample specific models

While the universal models trained on SEQC-II have shown to perform consistently better than other conventional somatic mutation detection schemes, here we explored whether sample-specific trained models could give us an additional accuracy boost, using nine replicate pairs across different SEQC-II datasets. For each sample, we trained NeuSomatic and NeuSomatic-S models using the *in silico* tumor-normal replicates prepared for that sample. We also trained a distinct model by combining 10% of training candidates used for the SEQC-WGS-Spike model and the training data derived for each sample. We compared these two sample-specific models with the universal SEQC-WGS-Spike model (**Suppl. Figure 6; Suppl. Table 9**). On average, the sample-specific models yielded ∼0.5% SNV and ∼5% INDEL F1-score improvements over the SEQC-WGS-Spike model for NeuSomatic. The library-prep and FFPE samples benefited the most from sample specific training. For instance, the library-prep sample prepared with Nextera Flex protocol with 1ng DNA amount gained 1.6% and 19.4% absolute F1-score improvements, respectively for SNVs and INDELs. Similarly, the 24h duration FFPE WGS sample with matched fresh normal gained 1.8 and 14.8 percentage points in F1-score improvement, respectively for SNVs and INDELs. For NeuSomatic-S, only INDELs seemed to benefit from sample-specific training.

### INDEL performance

We further evaluated the accuracies for detecting INDELs of different sizes across multiple datasets (**Figures 6a and 6b**). Although NeuSomatic did not explicitly use local assembly for INDEL detection, it still consistently surpassed other approaches including the assembly-based techniques like Lancet across a broad range of INDEL size, coverage, and tumor purity (**Figures 6a and 3d**). Overall, for insertions and deletions of more than 2 base pairs (bps), NeuSomatic, respectively, yielded 24.4% and 6.5% superiority in F1-score over Lancet, the best alternative conventional technique. Comparing INDEL accuracies across different NeuSomatic training approaches, the DREAM3 model suffered mainly from low insertion detection accuracy, whereas the SEQC-II trained models robustly identified both insertions and deletions of different sizes (**Figure 6b**).

**Figure 6.**
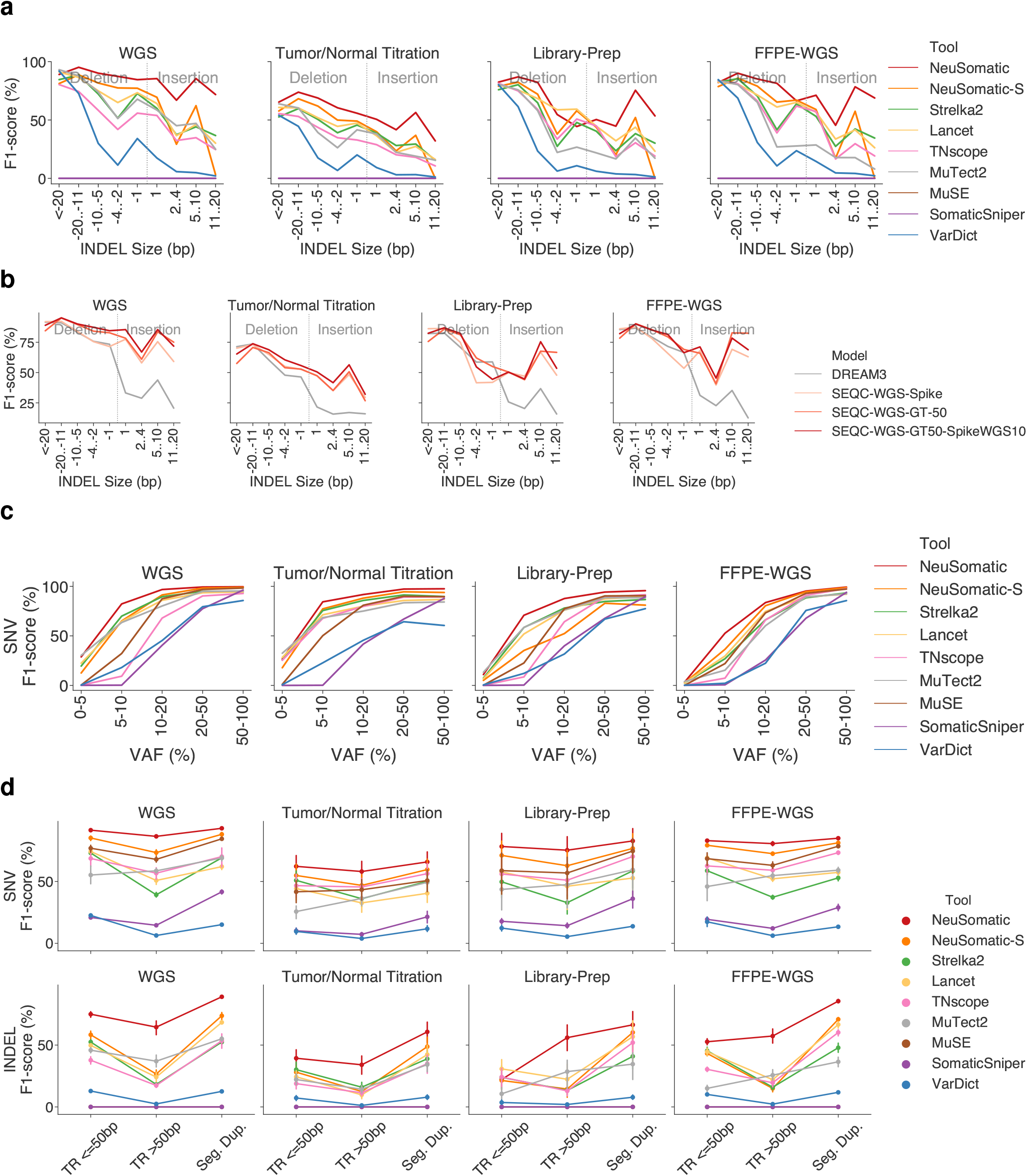
Performance analysis of different INDEL sizes, different VAF distributions, and on difficult regions. (a, b) Performance analysis for different INDEL sizes across SEQC-II datasets using different callers and different training approaches. Negative INDEL sizes reflect deletion. (c) Performance analysis for mutations with different VAF ranges across SEQC datasets using different callers. (d) Performance analysis of different somatic callers on difficult regions including tandem repeats (TR) of different sizes and segmental duplications.

### Performance analysis for different Variant Allele Frequencies (VAFs)

We analyzed the accuracy of different approaches across somatic mutation VAFs (**Figure 6c; Suppl. Figures 7-9**). On different datasets, we observe NeuSomatic show an added advantage over other schemes for VAFs less than 20% (**Figure 6c; Suppl. Figure 7**). We also observed that NeuSomatic has high robustness to VAF variations with consistent predictions for as low as 5% VAF. The SEQC-II trained models clearly performed better than the DREAM3 model for mutations with 5%-20% VAFs (**Suppl. Figure 8**). Similar robustness was observed for all coverage and tumor purity settings (**Suppl. Figure 9**).

### Performance on difficult genomic regions

To illustrate robustness to genomic context complexity, we evaluated the performance of different somatic calling schemes on difficult genomic regions including different length tandem repeats (TR) and segmental duplications (**Figure 6d; Suppl. Figure 10-13**). While many other schemes showed a significant drop in their performance on difficult genomic regions (**Suppl. Figure 10**), NeuSomatic remained highly robust and yielded even more significant improvement over other schemes compared to the whole genome analysis (**Figure 6d; Suppl. Figure 10**). On average, for these difficult genomic regions, NeuSomatic yielded more than 15% higher F1-scores than the best alternative conventional somatic caller for both SNVs and INDELs. Training NeuSomatic with SEQC-II samples resulted in large accuracy improvement on these difficult genomic regions, where the largest advantage was in tandem repeat regions using the models trained on ground truth mutations (**Suppl. Figures 11 and 12**). Similar performance on difficult genomic regions was observed across varying coverage and tumor purity (**Suppl. Figure 13**).

## Discussion

We explored the robustness of NeuSomatic’s deep learning framework in detecting somatic mutations across a diverse set of experimental settings seen in real WGS, WES, and AmpliSeq samples with different coverages, sample purities, and library preparations, on FFPE and fresh DNA sequenced with multiple platforms in six centers. Experiments confirmed the potential of NeuSomatic in capturing true somatic mutation signatures from the raw data to differentiate true calls from sequencing artifacts. NeuSomatic models trained on SEQC-II reference samples, both using spike-in mutations and the set of ground truth somatic mutations derived by SEQC-II studies, boosted the performance to achieve the highest accuracy. This analysis highlighted model-building best practice strategies, of utility in various scenarios. Compared with the baseline DREAM3 network models, the proposed models derived from SEQC-II reference samples were shown to reduce false positives for both SNVs and INDELs, improve insertion calling accuracies, and enhance somatic detection on difficult genomic regions by learning correct mutation signals.

While NeuSomatic remained robust to tumor contamination in the match normal, many somatic calling approaches like MuTect2, Lancet, MuSE, and TNscope were highly impacted, because they rejected true somatic mutations that had corresponding reads in normal. Thus, their recall rate dropped significantly. NeuSomatic models trained on WGS performed equally well on targeted sequencing data. On the other hand, TNscope employs a machine learning (random forest) model, while Lancet uses localized colored de Bruijn graphs approach to detect mutations. Both tools were designed for genome-wide somatic mutation detection and thus are not suitable for targeted sequencing (typically covers less than 1 MB of the genome).

Analyzing prediction accuracies across different specifications including mutation type, INDEL length, VAF, and genomic region revealed that NeuSomatic yielded the largest improvements over other schemes in challenging situations such as low coverage, less than 20% VAF, difficult genomic regions like tandem repeats, INDELS more than 2bp long, FFPE with DNA damage, or contaminated match normal. As a result of confusing sequencing or sample-specific artifacts with the true somatic mutations, conventional schemes had many false positives (and thus lower precision), often seen in low-complexity genomic regions (**Figure 6d; Suppl. Figure 10**). However, by training on WGS samples from multiple platforms and multiple centers, NeuSomatic learned error-prone genomic contexts and thus consistently enhanced accuracy across different conditions including difficult-to-call low-complexity genomic contexts. Similarly, analysis of missed calls by different approaches revealed that most of the private false negative calls by conventional schemes like Strelka2, which were correctly predicted by NeuSomatic, had low VAF (**Suppl. Figure 14**). This revealed the inferiority of other schemes in detecting low VAF mutations. NeuSomatic, on the other hand, more accurately distinguished challenging low VAF calls from artifacts by learning the correct signature from raw data.

## Methods

### SEQC-II tumor-normal sequencing data and ground truth

We used the SEQC-II reference matched samples, a human triple-negative breast cancer cell line (HCC1395) and a matched B lymphocyte derived normal cell line (HCC1395BL) in our analysis. Detailed sample information can be found in the SEQC-II reference samples manuscripts^12,13^. The SEQC-II somatic mutation working group has established a gold standard truth set of somatic mutations for these samples^13^ (**Suppl. Figure 1**). The truth set was defined using multiple tumor-normal sequencing replicates from different sequencing centers, and orthogonal mutation detection bioinformatics pipelines. To assign confidence scores to the mutation calls and minimized platform, center, and pipeline specific biases, the SomaticSeq^10^ machine learning framework was implemented to train a set of classifiers on *in silico* tumor-normal replicate pairs generated by spiking synthetic mutations into an HCC1395BL alignment file, matched with another HCC1395BL for each replicate pair. Using these classifiers, mutation calls were categorized into four confidence-levels (HighConf, MedConf, LowConf, and Unclassified) based on cross-aligner and cross-sequencing-center reproducibility. HighConf and MedConf calls were grouped together as the “truth set” of somatic mutations (v1.0), which contains a total of 39,536 SNVs and 2,020 INDELs. The truth set for somatic mutations in HCC1395, is available for community use on NCBI’s ftp site (ftp://ftp-trace.ncbi.nlm.nih.gov/seqc/ftp/Somatic_Mutation_WG/).

All sequencing data used in this study are publicly available through NCBI’s SRA database (SRP162370). For all samples, the FastQ files were first trimmed using Trimmomatic^15^, and then aligned with BWA-MEM (v0.7.15)^16^ followed by Picard MarkDuplicates (https://broadinstitute.github.io/picard).

### Training datasets and models

Different training datasets were used to derive effective NeuSomatic CNN models using both *in silico* tumor replicates, prepared with synthetic spike-in mutations, as well as real tumor replicates with known high confidence truth set of somatic mutations (**Suppl. Table 1**):

#### DREAM3 model

As the baseline WGS model we employed the DREAM3 model, developed recently^11^ by training on ICGC-TCGA DREAM Challenge Stage 3 data^14^. The Stage 3 dataset consists of a normal sample and a tumor sample constructed by computationally spiking 7,903 SNVs and 7,604 INDELs mutations into a healthy genome of the same normal sample with three different AFs of 50%, 33%, and 20% to create synthetic but realistic tumoral normal pairs. An additional ∼95K SNVs and ∼95K INDELs were spike-in into the tumor sample^11^ using BAMSurgeon^17^ with similar AF distributions to the original DREAM data for better network training. This model was trained by combining training data (in 50% of the genome) from five different tumor-normal purity setting of 100T:100N, 50T:100N, 70T:95N, 50T:95N, and 25T:95N^11^. NeuSomatic and NeuSomatic-S were trained on ∼29.2M candidate mutations identified in these five replicate pairs, which includes ∼450K candidates with somatic SNV/INDEL labels and ∼28.7M candidates, labeled as non-somatic.

#### TCGA model

As the baseline WES model, we used the TCGA model developed recently^11^ by training on a set of 12 TCGA samples^18^. The tumor and normal alignments for each of these samples were mixed and split into two equal alignments. One alignment was treated as the normal replicate and the other alignment was used to spike in mutations to construct the tumor replicate. For each sample, ∼88K SNVs and ∼44K INDELs were spiked in to generate a synthetic tumor sample for training. NeuSomatic and NeuSomatic-S were trained on ∼5.9M candidate mutations identified in these 12 replicate pairs, which includes ∼1.5M candidates with somatic SNV/INDEL labels and ∼4.4M candidates, labeled as non-somatic.

#### SEQC-WGS-Spike model

To prepare the training data used to build this model, we employed BAMSurgeon to construct a synthetic tumor by spiking *in silico* mutations in one of the HCC1395BL replicates and pairing that with distinct HCC1395BL replicates as normal match. Using this approach, we prepared ten *in silico* tumor-normal pairs. Eight of the ten replicate pairs were selected from WGS replicates sequenced in four different sites with 40×-95× mean coverage. The other two replicate pairs were created by merging multiple NovaSeq sequencing replicates from Illumina to obtain *in silico* ∼220× tumor sample coverage and ∼170× normal matched sample coverage. In each *in silico* tumor we spiked in ∼92 K SNVs and ∼22 K INDELs. The AFs of the spike-in mutations were randomly selected from a beta distribution (with shape parameters ***α***=2 and ***β***=5). For each of the ten replicate pairs we also constructed an impure normal by mixing 95% normal and 5% tumor reads. So, in total we used 20 *in silico* tumor-normal pairs to train the SEQC-WGS-Spike model. We then trained NeuSomatic and NeuSomatic-S on ∼272M candidate mutations identified in these 20 replicate pairs, which included ∼2M candidates with somatic SNV/INDEL labels and ∼270M candidates, labeled as non-somatic.

#### SEQC-WGS-GT-50 model

This model was constructed using the real WGS tumor-normal replicates accompanied by the SEQC-II HighConf truth set as the ground truth somatic mutations. We used eight WGS tumor-normal replicates as the base samples to train this model. The first seven WGS replicate pairs were from six different sequencing centers on HiSeq and NovaSeq platforms with mean coverage of 40×-95×, and the last one was constructed by combining 9 NovaSeq sequencing replicates from Illumina to get a replicate pair with ∼390× coverage. For each of these eight replicate pairs, we constructed two other purity variations, one with 95% normal purity by mixing 95% normal and 5% tumor reads, and the other with 10% tumor purity by mixing 10% tumor and 90% normal reads. So, for each of the replicate pairs we had a version with 100% pure tumor and normal, a version with 100% pure tumor matched with 95% pure normal, and a version with 10% pure tumor matched with 100% pure normal. Thus, in total, we used 24 tumor-normal replicates to train the SEQC-WGS-GT-50 model. To have unbiased evaluation, we only used 50% of the genome to train this model and kept the other 50% for evaluation. To prepare the training and evaluation regions, we split the genome to small regions of size ∼92K bps, and randomly select half of the fractured regions for training and the other half for evaluation. We excluded a 5-bases padded region around each of the grey-zone mutations, including calls with LowConf label in the super-set of SEQC-II calls, and those with Unclassified label in the super-set of SEQC-II calls with more than 30% VAF, from the training region. We then trained NeuSomatic and NeuSomatic-S on ∼137M candidate mutations identified in these 24 replicate pairs, which included ∼416K candidates with somatic SNV/INDEL labels and ∼136.5M candidates, labeled as non-somatic.

#### SEQC-WGS-GT-ALL model

This model was prepared similar to the SEQC-WGS-GT-50 model but using all HighConf ground truth somatic mutations over the whole genome. So, it cannot be used to evaluate WGS samples which were used for training purposes and also may be biased towards truth set mutations given all have been used for training. This model is still beneficial for performance analysis on other datasets or non-SEQC-II samples. We trained this model for NeuSomatic and NeuSomatic-S on ∼272M candidate mutations identified in the 24 replicate pairs used for SEQC-WGS-GT-50 model, which included ∼839K candidates with somatic SNV/INDEL labels and ∼271M candidates, labeled as non-somatic.

#### SEQC-WGS-GT50-SpikeWGS10 model

The training data for this model was prepared by combining 10% of training candidates used for SEQC-WGS-Spike model and all candidates from SEQC-WGS-GT-50. This combination takes advantage of both a high number of spike-in mutations and the real somatic mutation characteristics seen in real tumor-normal data. We trained NeuSomatic and NeuSomatic-S on the combined set of 164M candidate mutations, including ∼574K candidates with somatic SNV/INDEL labels and ∼163.5M candidates, labeled as non-somatic.

#### SEQC-WES-Spike model

Similar to the SEQC-WGS-Spike model, we used BAMSurgeon to construct 7 *in silico* tumor-normal WES replicates to train this model. The replicate pairs were selected from the WES dataset sequenced in four different sites with 60×-550× mean coverage. In each *in silico* tumor we spiked in ∼97 K SNVs and ∼19 K INDELs. The AF of the spike-in mutations were randomly selected from a beta distribution (with shape parameters ***α***=2 and ***β***=5). We then trained NeuSomatic and NeuSomatic-S on ∼3.7M candidate mutations identified in these 7 replicate pairs, which included ∼755K candidates with somatic SNV/INDEL labels and ∼3M candidates, labeled as non-somatic.

#### SEQC-FFPE-Spike model

Similar to the SEQC-WGS-Spike model, we used BAMSurgeon to construct 8 *in silico* tumor-normal WGS FFPE replicates to train this model. The replicate pairs were selected from the FFPE dataset sequenced at two different sites with four different preparation times. In each *in silico* tumor we spiked in ∼92 K SNVs and ∼22 K INDELs. The AF of the spike-in mutations were randomly selected from a beta distribution (with shape parameters ***α***=2 and ***β***=5). We also matched each *in silico* tumor with a fresh WGS replicate to include FFPE tumor versus fresh normal scenarios in our training. So, in total we used 16 *in silico* tumor-normal pairs to train the SEQC-FFPE-Spike model. We trained NeuSomatic and NeuSomatic-S on ∼191M candidate mutations identified in these 7 replicate pairs, which included ∼1.7M candidates with somatic SNV/INDEL labels and ∼190M candidates, labeled as non-somatic.

#### SEQC-FFPE-WES-Spike model

Similar to other spike-in models, we used BAMSurgeon to construct 7 *in silico* tumor-normal WES FFPE replicates to train this model. The replicate pairs were selected from the WES FFPE dataset sequenced in two different sites and prepared across four varying time intervals. Since we don’t have two normal replicates with the same FFPE preparation time and sequencing site for this dataset, we mixed the tumor and normal alignments for each replicate pair and split the mixture into two equal alignments. We then treat one alignment as the normal replicate and spike-in mutations in the other to construct the tumor replicate. In each *in silico* tumor we spiked in ∼90 K SNVs and ∼19 K INDELs. The AF of the spike-in mutations were randomly selected from a beta distribution (with shape parameters ***α***=2 and ***β***=5). We also matched each *in silico* tumor with a fresh WES replicate to include FFPE tumor versus fresh normal scenarios in our training. So, in total, we used 14 *in silico* tumor-normal pairs to train the SEQC-FFPE-WES-Spike model. We trained NeuSomatic and NeuSomatic-S on ∼9.6M candidate mutations identified in these 7 replicate pairs, which included ∼1.4M candidates with somatic SNV/INDEL labels and ∼8.2M candidates, labeled as non-somatic.

#### SEQC-WES-SpikeWGS10, SEQC-FFPE-SpikeWGS10, SEQC-FFPE-WES-SpikeWGS10 models

The training data for these models was prepared by combining 10% of training candidates used for SEQC-WGS-Spike model and respectively all candidates from SEQC-WES-Spike, SEQC-FFPE-Spike, and SEQC-FFPE-WES-Spike. This combination takes advantage of both the high number of spike-in WGS mutations, as well as the sample biases for WES and FFPE samples.

#### Sample-specific models

In addition to the above general-purpose models, for a set of nine samples across multiple data types, we derived sample-specific models. The selected samples included a WGS sample sequenced by a NovaSeq instrument, a WES sample, a sample prepared with a Nextera Flex library-prep kit with 1ng DNA amount, a 30× WGS sample with 50% tumor purity, a 100× WGS sample with 10% tumor purity, two WGS and two WES FFPE samples each treated with formalin for 24h and matched with fresh/FFPE normal samples. For each sample, we prepared the *in silico* tumor using random spikes. For the 10% tumor sample, the AF of the spike-in mutations were randomly selected from a beta distribution (with shape parameters ***α***=2 and ***β***=18). For the other samples, a beta distribution (with shape parameters ***α***=2 and ***β***=5) was used to select the AFs. For each sample, we then trained NeuSomatic and NeuSomatic-S models using the *in silico* tumor-normal replicates. In addition, for each sample we trained a distinct model by combining 10% of training candidates used for SEQC-WGS-Spike model and the training data derived for that sample.

### Somatic mutation detection algorithms

In addition to NeuSomatic, we used seven somatic mutation callers, MuTect2 (4.beta.6)^2^, SomaticSniper (1.0.5.0)^4^, Lancet (1.0.7)^8^, Strelka2 (2.8.4)^5^, TNscope (201711.03)^7^, MuSE (v1.0rc)^3^, and VarDict (v1.5.1)^6^ and ran each of them using the default parameters or parameters recommended by the user’s manual. For SomaticSniper, we used “-q 1 -Q 15 -s 1e-05”. For TNscope (201711.03), we used the version implemented in Seven Bridges’s CGC with the following command, “sentieon driver -i $tumor_bam -i $normal_bam -r $ref --algo TNscope --tumor_sample $tumor_sample_name --normal_sample $normal_sample_name -d $dbsnp $output_vcf”. For Lancet, we used “--cov-thr 10 --cov-ratio 0.005 --max-indel-len 50 -e 0.005”. The high confidence outputs flagged as “PASS” in the resulting VCF files were applied to our comparison analysis. For VarDict we also restricted to calls with “Somatic” status. Results from each caller used for comparison were all mutation candidates that users would otherwise consider as “real” mutations detected by this caller.

We used NeuSomatic (0.1.4) in both ensemble and standalone mode. For ensemble mode, in addition to the candidate variants identified by “alignment scanning” step of NeuSomatic, we also incorporated calls from five individual callers (MuTect2, SomaticSniper, Strelka2, MuSE, and VarDict) and represented their occurrence as additional input channels for each candidate variant. For NeuSomatic and NeuSomatic-S, we used “--scan_maf 0.01--min_mapq 10 --snp_min_af 0.03 --snp_min_bq 15 --snp_min_ao 3 --ins_min_af 0.02 --del_min_af 0.02 --num_threads 28 --scan_window_size 500 --max_dp 100000” during preprocessing step. For training, we used “--coverage_thr 300 --batch_size 8000”.

### Difficult genomic regions

We used a set of difficult genomic regions derived by the Genome-in-a-Bottle consortium^19^. These regions included tandem repeats, in two different categories of less than 50bp and larger than 50bp repeats, and segmental duplication regions. We evaluated different techniques on these regions to illustrate robustness to genomic context complexity.

### Analysis of false negative calls

To better understand the difference in performance of different techniques, we performed a set of pairwise comparison between the VAF distribution of the false negatives SNVs of NeuSomatic against other schemes on WGS dataset (**Suppl. Figure 14**). In each X-vs-Y analysis, we identified X_FN_, the set of ground truth SNVs which were missed by algorithm X (False Negative in X) in at least 11 out of 21 WGS replicates. Similarly, we identified Y_FN_, the set of ground truth SNVs which were missed by algorithm Y (False Negative in Y) in at least 11 out of 21 WGS replicates. We then computed the private false negatives of X, defined as X_FN_ / Y_FN_, and the private false negatives of Y, defined as Y_FN_ / X_FN_. For somatic mutations in each of those sets we then computed the distribution of VAF of mutations.

### Evaluation process

For a fair comparison, we evaluated all the models and somatic mutation algorithms on the 50% hold-out genomic region which was not used for training the SEQC-WGS-GT-50 model. This ∼1.4GB region contained ∼21K SNVs and ∼1.3K INDELs from the SEQC-II truth set for HCC1395 which were not used for training any of the NeuSomatic models except for SEQC-WGS-GT-ALL model.

Calls labeled as HighConf and MedConf by SEQC-II consortium for HCC1395 were grouped together as the “truth set” of somatic mutations used herein. We employed this truth set to compute the true positives and false negatives across all replicates of HCC1395 for all pipelines. As recommended by SEQC-II consortium, we also blacklisted LowConf calls as they have low validation rates. Since this truth set provided by the consortium had a VAF detection limit of 5% and depth detection limit of 50×, for calls at higher coverages data or lower VAFs, we cannot ascertain their real somatic status. Thus, for private calls reported by any pipeline that was not in the truth set, we excluded calls which were deemed to be ambiguous for evaluation. In summary, for a tumor replicate in-hand that had a mean coverage of *C* and a tumor purity of *P*, if a pipeline had reported a private somatic mutation for this replicate (which was not in the truth set) that has *d* number of supporting reads, we only labeled this call as false positive if its projected number of supporting reads at 100% purity and 50x coverage was ≥ 3. In other words, this call was false positive if *d* ≥ 3*CP*/50; otherwise we exclude this call from evaluation.

For WES and AmpliSeq data since the number of true indels in the evaluation region was very limited, we only reported SNVs evaluations.

## Acknowledgements

We thank Dr. Rebecca Kusko of the Immuneering Corporation for manuscript polishing and Dr. Jun Ye of Sentieon for providing the Sentieon software package. We also appreciate Dr. Laufey Amundadottir of the Division of Cancer Epidemiology and Genetics, National Cancer Institute (NCI), National Institutes of Health (NIH), for the sponsorship and the usage of the NIH Biowulf cluster; Seven Bridges for providing storage and computational support on the Cancer Genomic Cloud (CGC). The CGC has been funded in whole or in part with Federal funds from the National Cancer Institute, National Institutes of Health, Contract No. HHSN261201400008C and ID/IQ Agreement No. 17X146 under Contract No. HHSN261201500003I. This work also used the computational resources of the NIH Biowulf cluster (http://hpc.nih.gov). Original data was also backed up on the servers provided by Center for Biomedical Informatics and Information Technology (CBIIT), NCI.

## Disclaimer

This is a research study, not intended to guide clinical applications. The views presented in this article do not necessarily reflect current or future opinion or policy of the US Food and Drug Administration. Any mention of commercial products is for clarification and not intended as endorsement.

## Author contributions

Study conceived and designed by: W.X., S.M.E.S., L.T.F, and M.M

Data analysis: S.M.E.S., W.X., L.T.F, and M.M

Manuscript writing: S.M.E.S., W.X., L.T.F, M.M, and H.H

