## Supplementary Information for "Robust Cancer Mutation Detection with Deep Learning Models Derived from Tumor-Normal Sequencing Data"

### Supplementary Figures

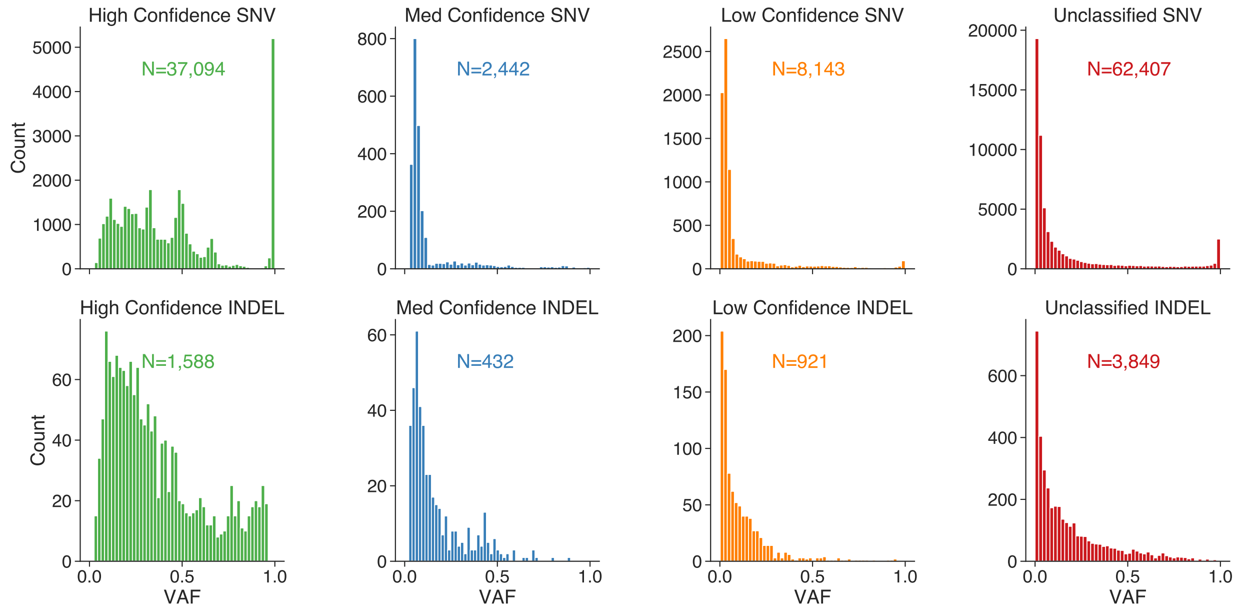


Supplementary Figure 1: VAF distribution of the ground truth SNV and INDEL somatic mutations in the super set of calls for HCC1395 classified by SEQC-II consortium to four confidence levels (High, Med, Low, and Unclassified). High and medium confidence calls are grouped together as the “truth set” of somatic mutations.

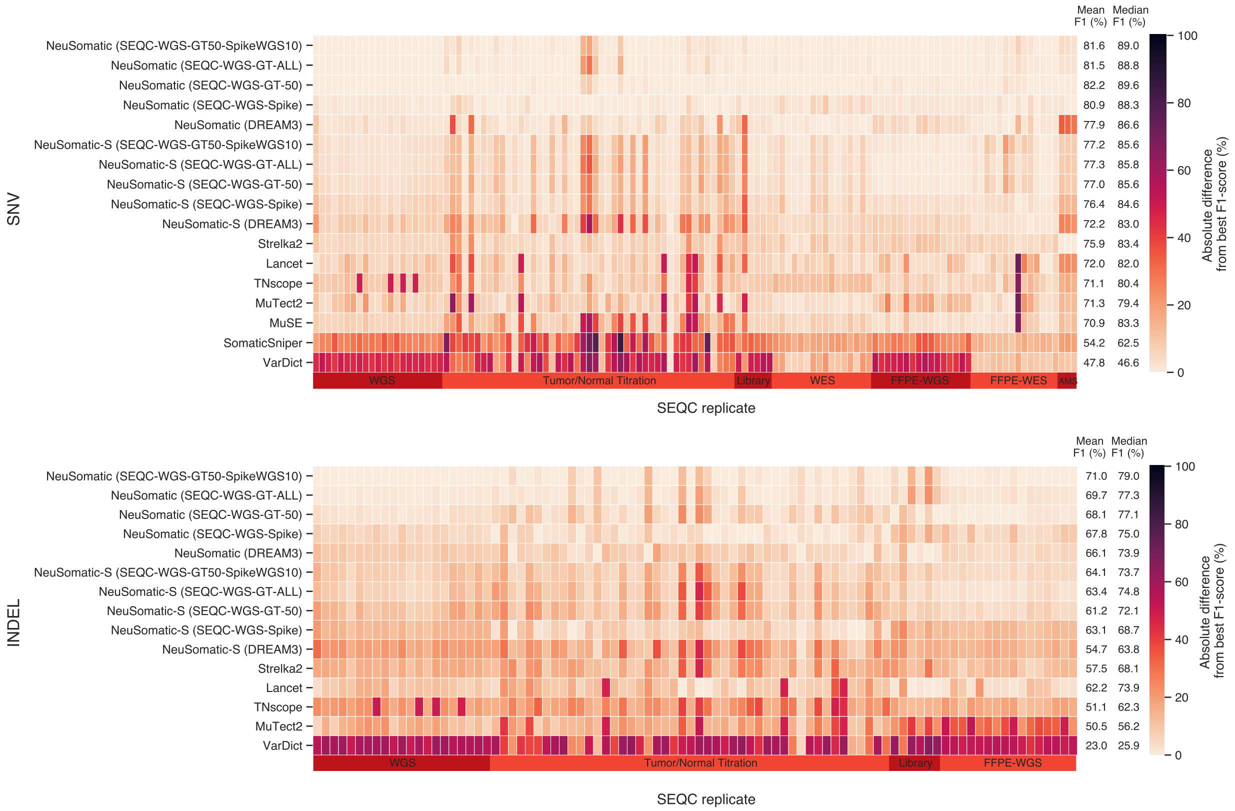


Supplementary Figure 2: Performance comparison of different techniques across 123 replicates in six datasets for SNVs and INDELs. For each replicate the best F1-score was computed across different approaches. The heatmaps illustrates the absolute difference between the F1-score of any of the somatic mutation detection approaches to the best F1-sScore. The mean F1-score is shown for each approach across 123 replicates.

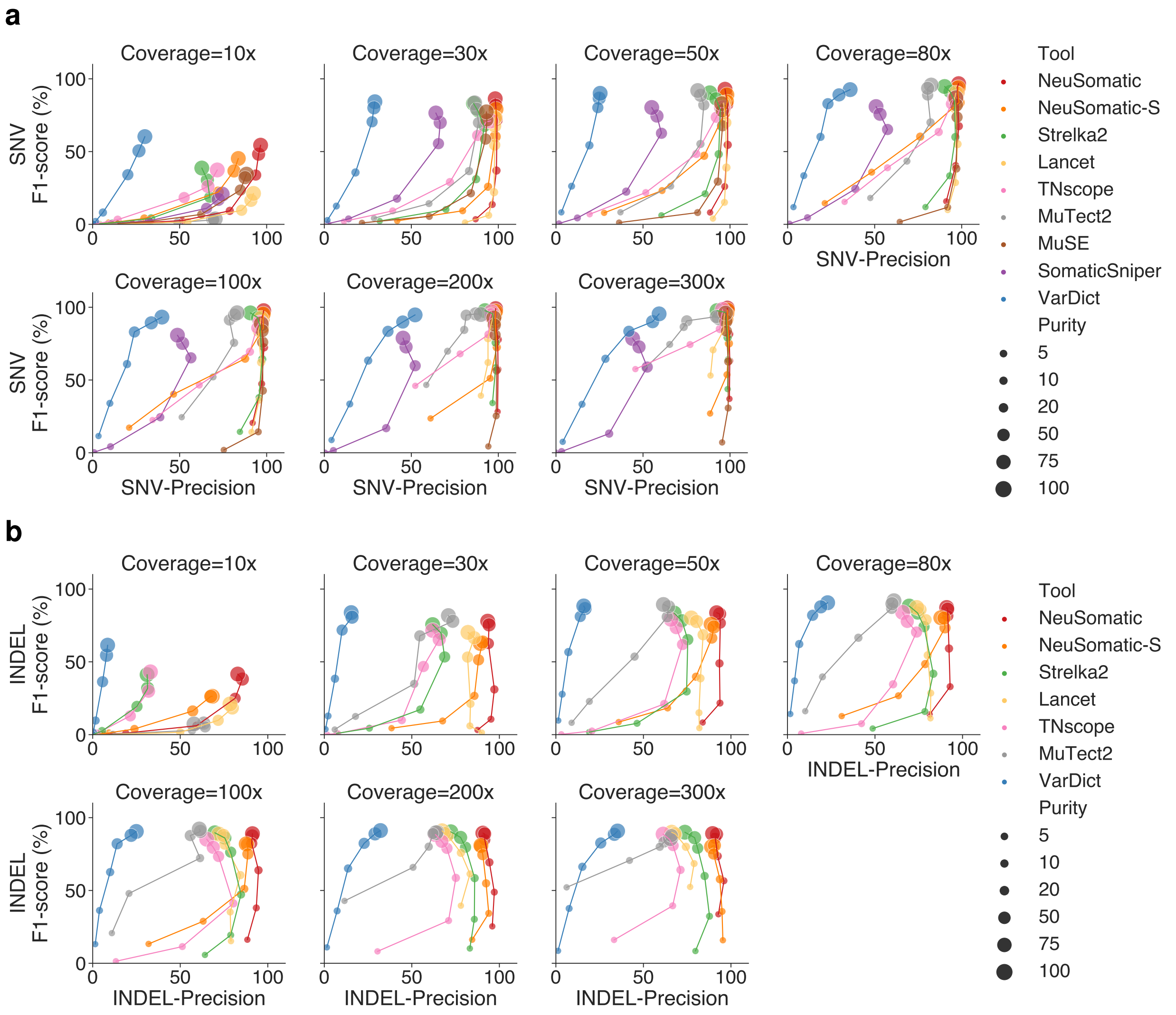


Supplementary Figure 3: Precision-Recall analysis on Tumor-purity dataset. (a) SNV and (b) INDEL accuracies are compared for different somatic mutation callers across different coverages (10x-300x) and tumor purities (5%-100%).

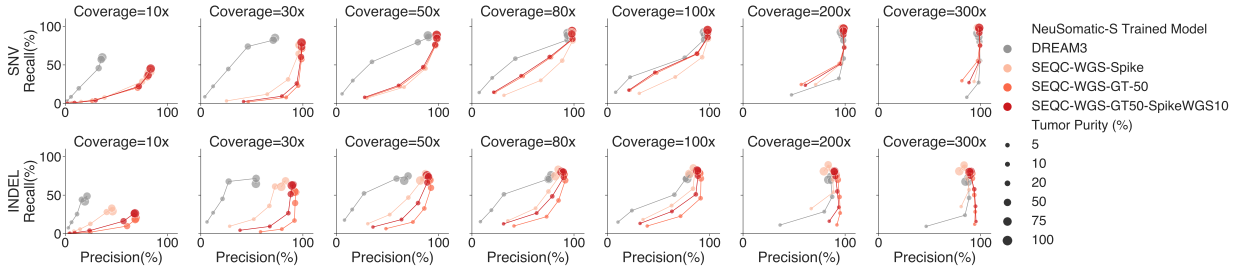


Supplementary Figure 4: Precision-Recall comparison for various NeuSomatic-S trained models on the tumor-normal titration dataset.

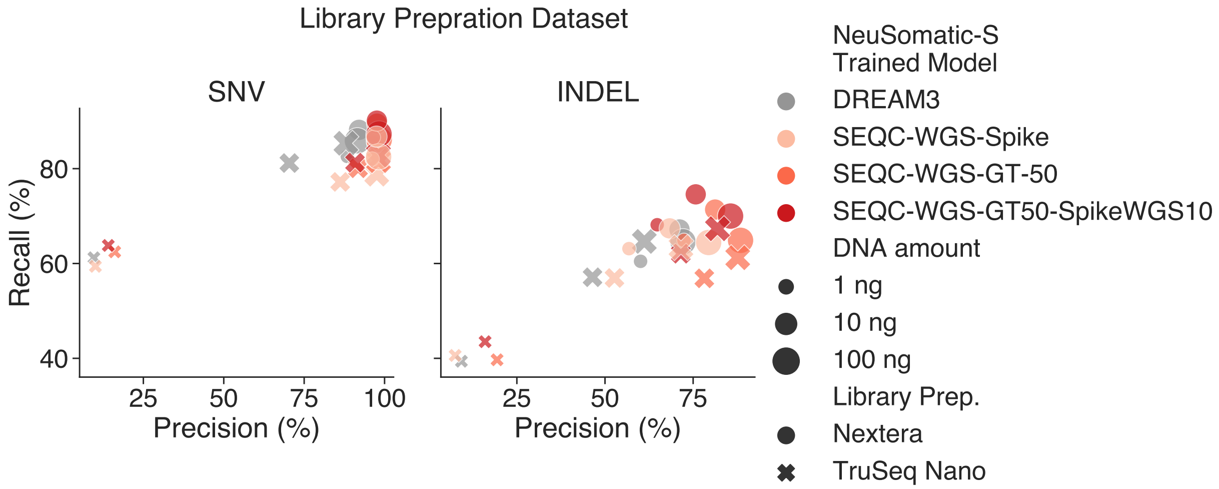


Supplementary Figure 5: Precision-Recall comparison for various NeuSomatic-S trained models on the Library preparation dataset.

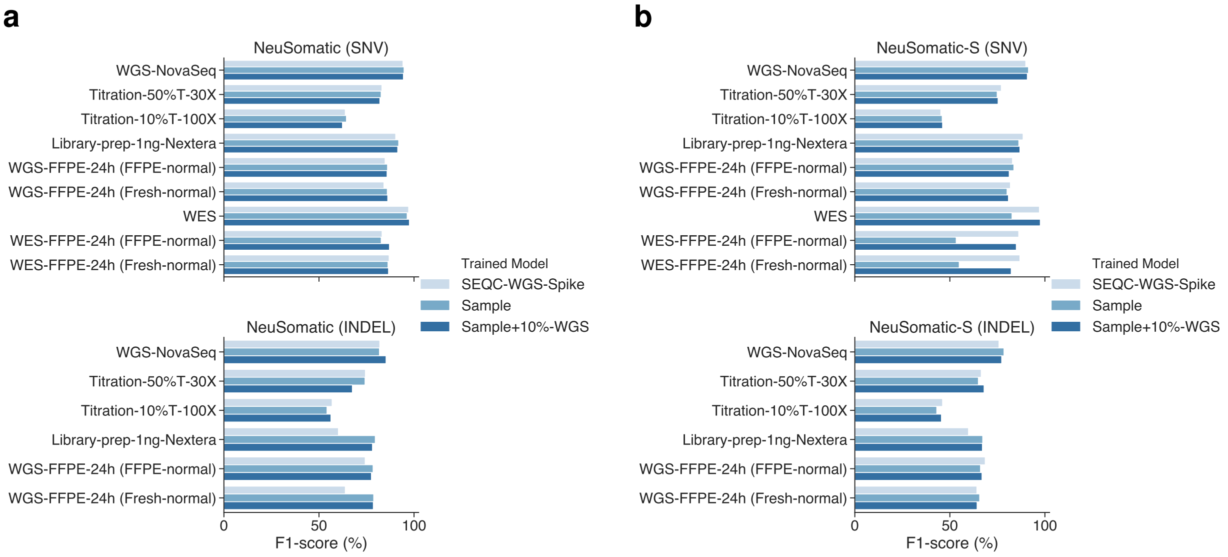


Supplementary Figure 6: Analyzing the impact of training on the target sample for nine replicates from different datasets using (a) NeuSomatic and (b) NeuSomatic-S. For each sample, two sample specific models were used: one only trained on the target sample, and the other with an additional 10% of the training data from SEQC-WGS-Spike model.

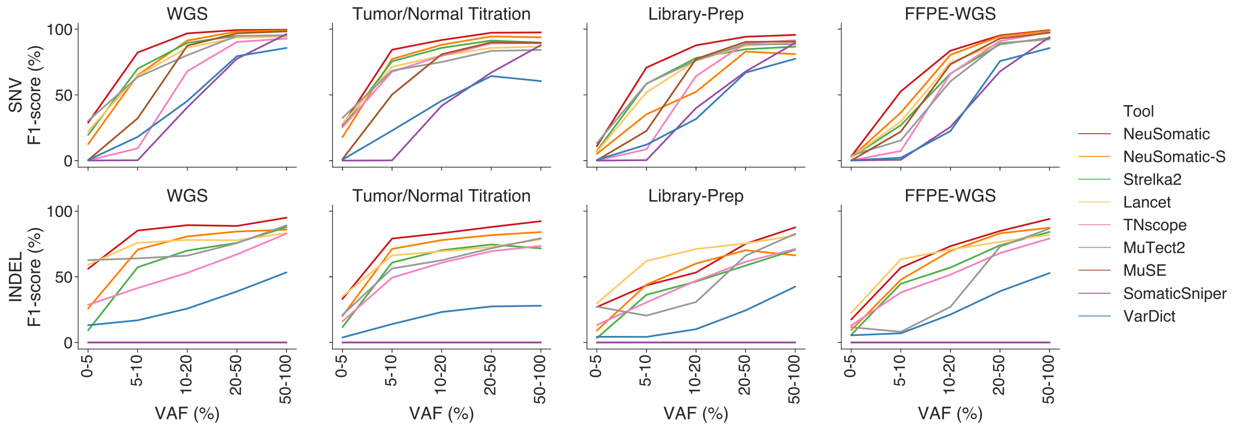


Supplementary Figure 7: Performance analysis for mutations with different VAF ranges across SEQC datasets using different callers.

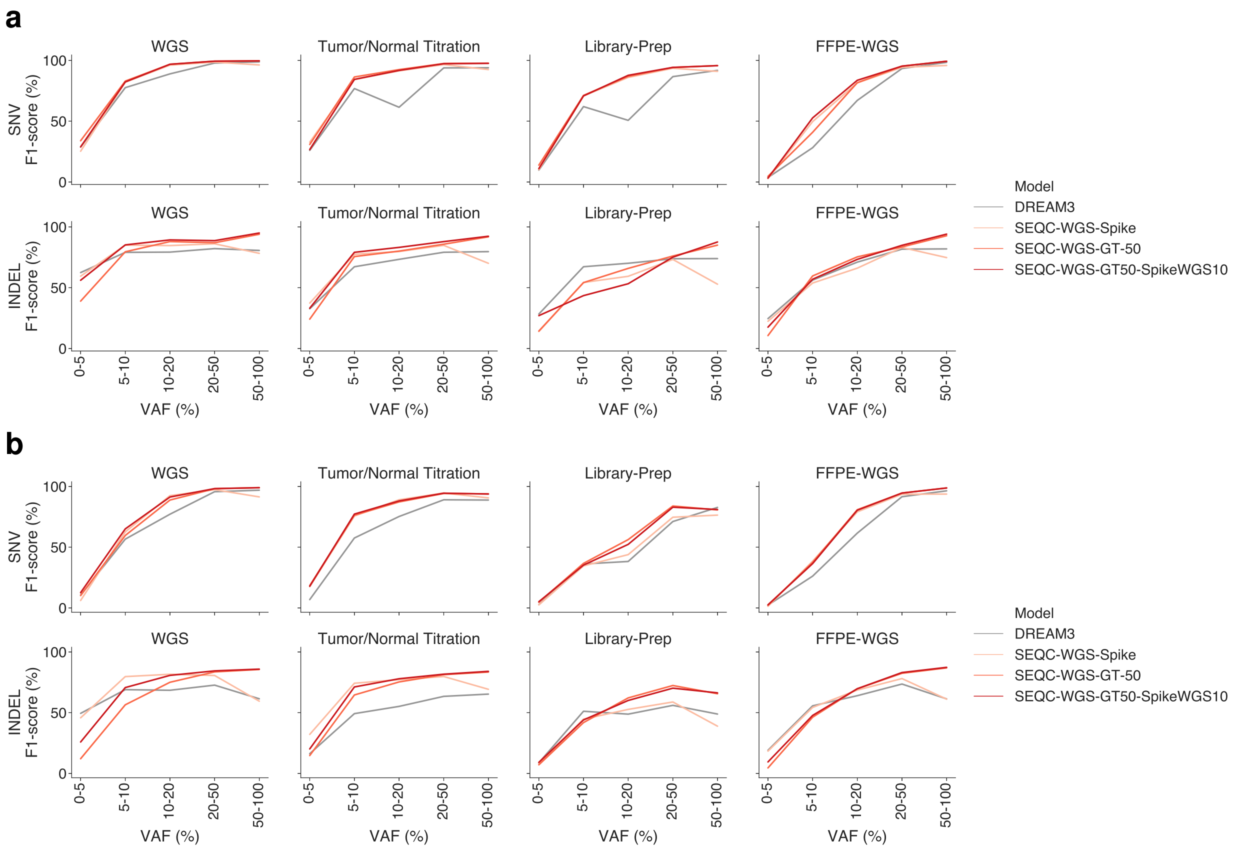


Supplementary Figure 8: Performance analysis for mutations with different VAF ranges across SEQC datasets using different training approaches for (a) NeuSomatic and (b) NeuSomatic-S.

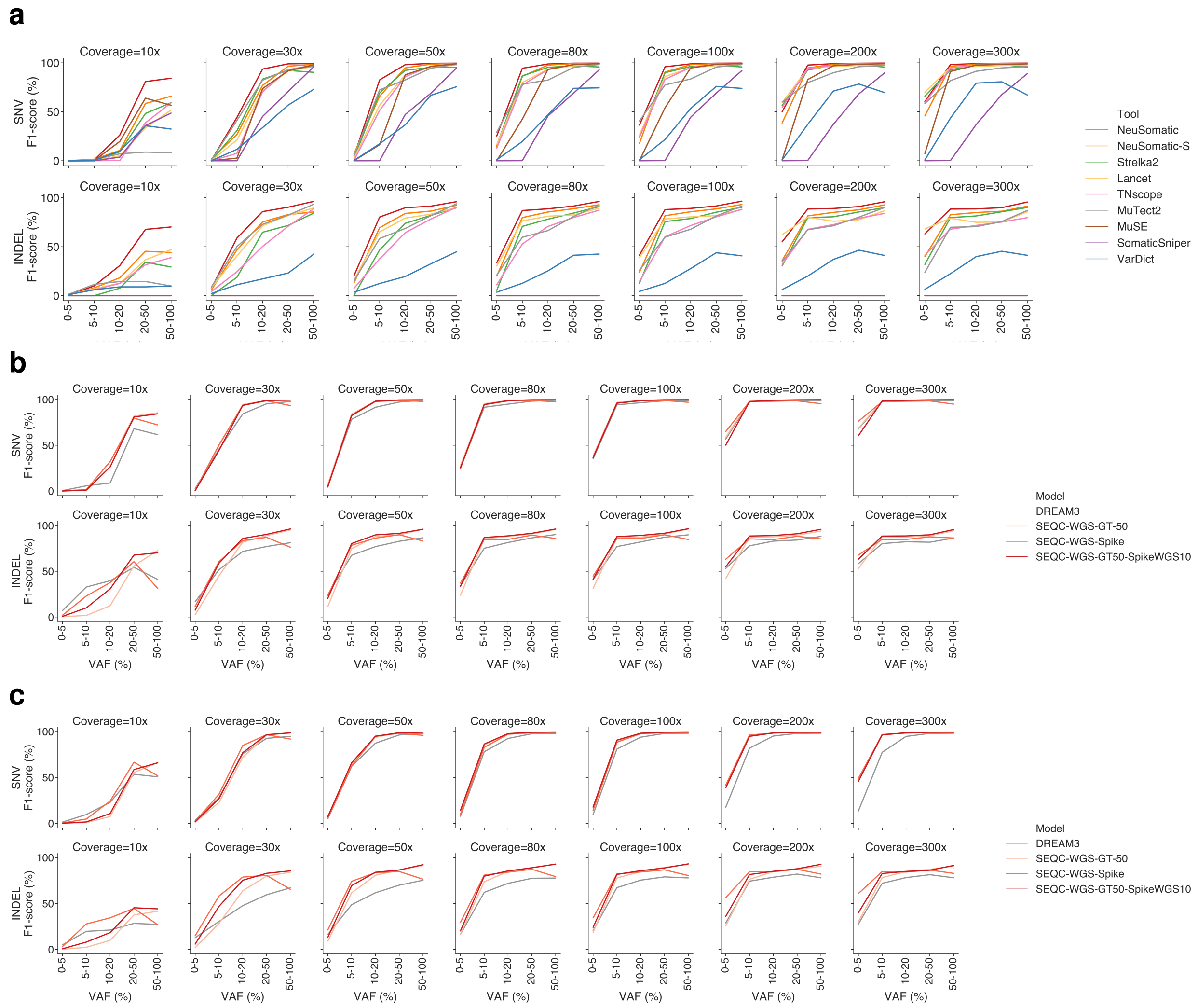


Supplementary Figure 9: Performance analysis for mutations with different VAF ranges on Tumor/Normal titration dataset using different (a) callers and (b, c) training approaches for (b) NeuSomatic and (c) NeuSomatic-S.

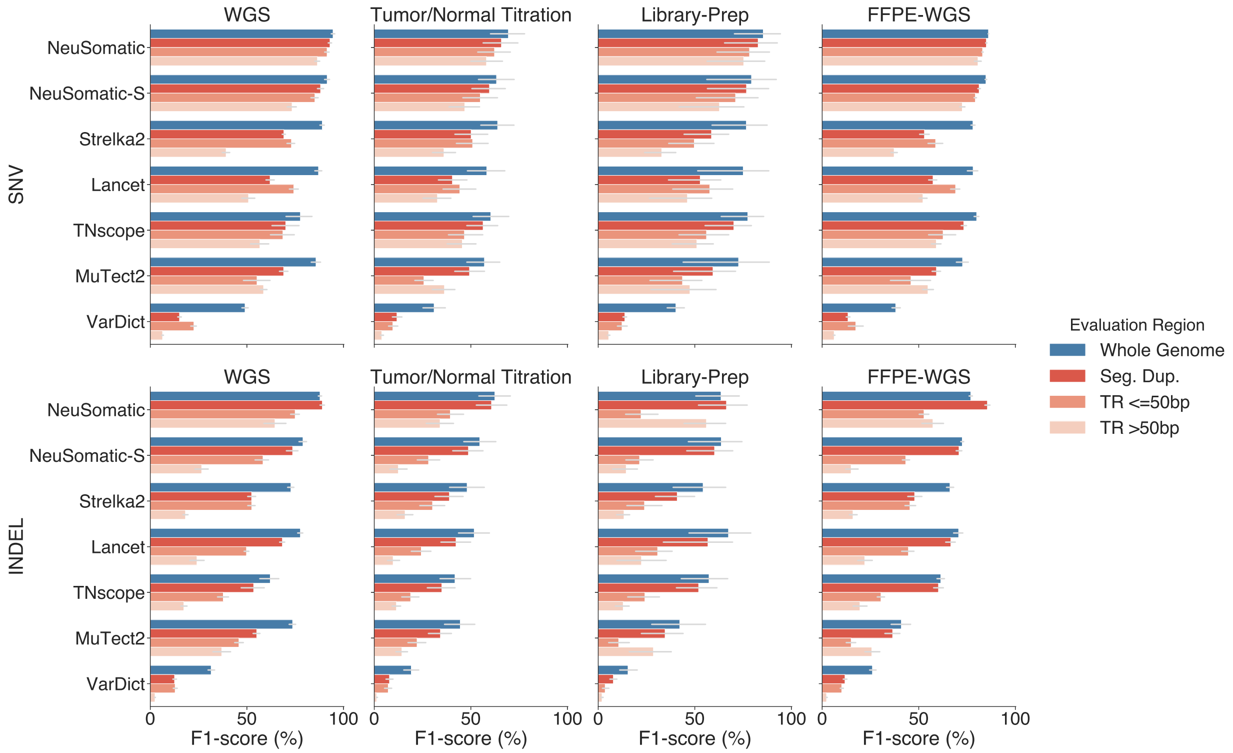


Supplementary Figure 10: Performance comparison on whole genome versus the difficult genomic regions including tandem repeats (TR) of different sizes and segmental duplications using different callers.

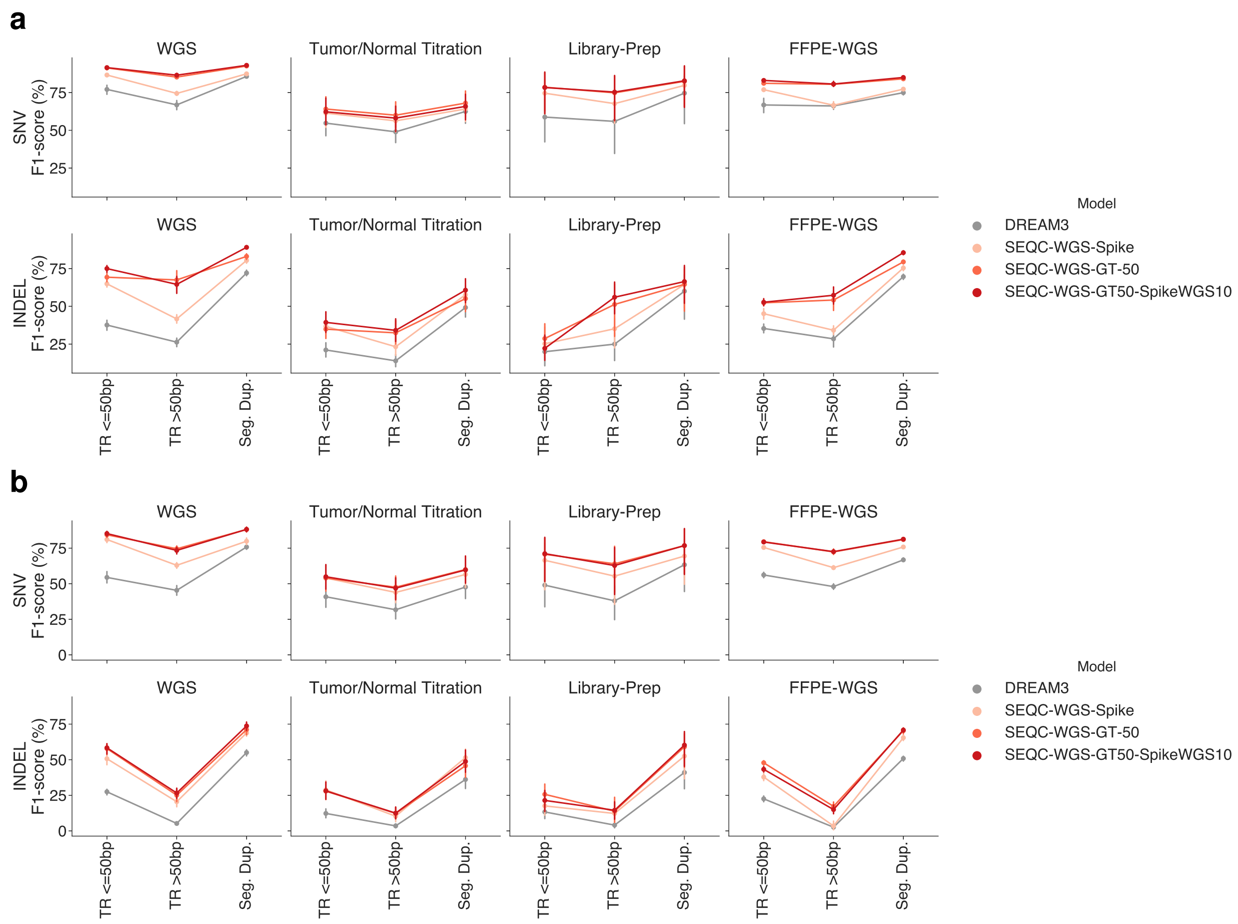


Supplementary Figure 11: Performance analysis of different training approaches on difficult regions including tandem repeats (TR) of different sizes and segmental duplications for (a) NeuSomatic, (b) NeuSomatic-S.

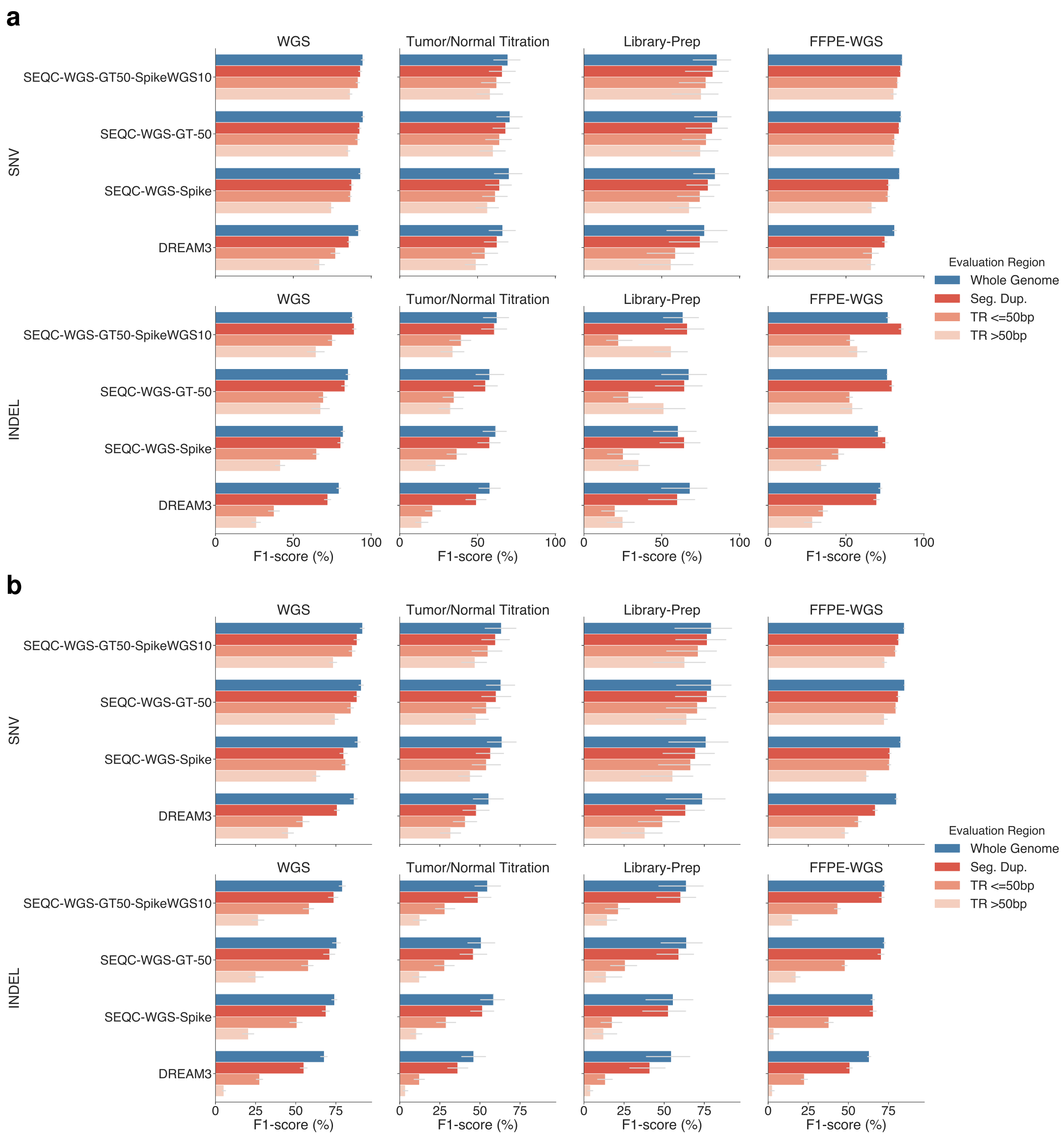


Supplementary Figure 12: Performance comparison on whole genome versus the difficult genomic regions including tandem repeats (TR) of different sizes and segmental duplications using different training approaches for (b) NeuSomatic and (c) NeuSomatic-S.

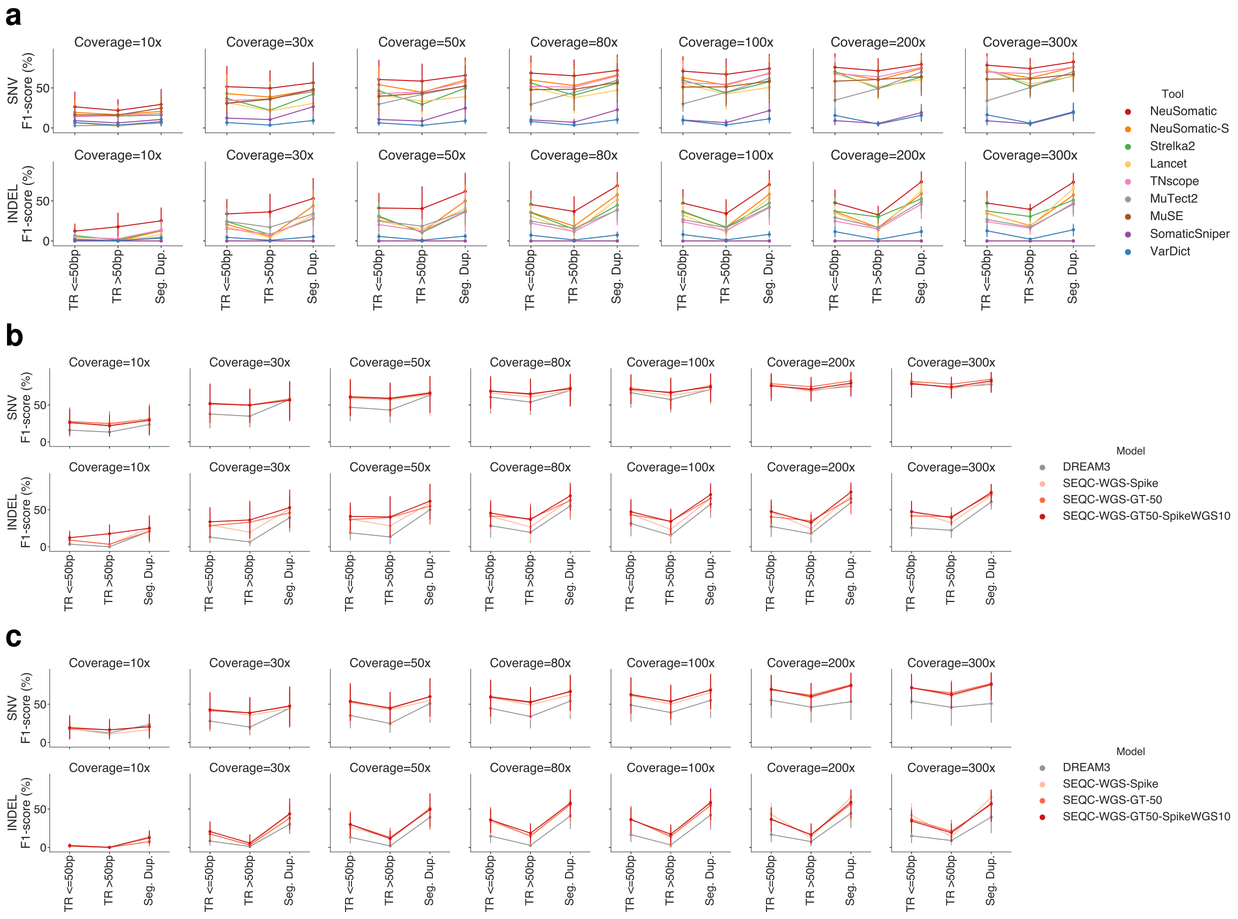


Supplementary Figure 13: Performance analysis on difficult regions including tandem repeats (TR) of different sizes and segmental duplications on Tumor/Normal titration dataset using different (a) callers and (b, c) training approaches for (b) NeuSomatic and (c) NeuSomatic-S.

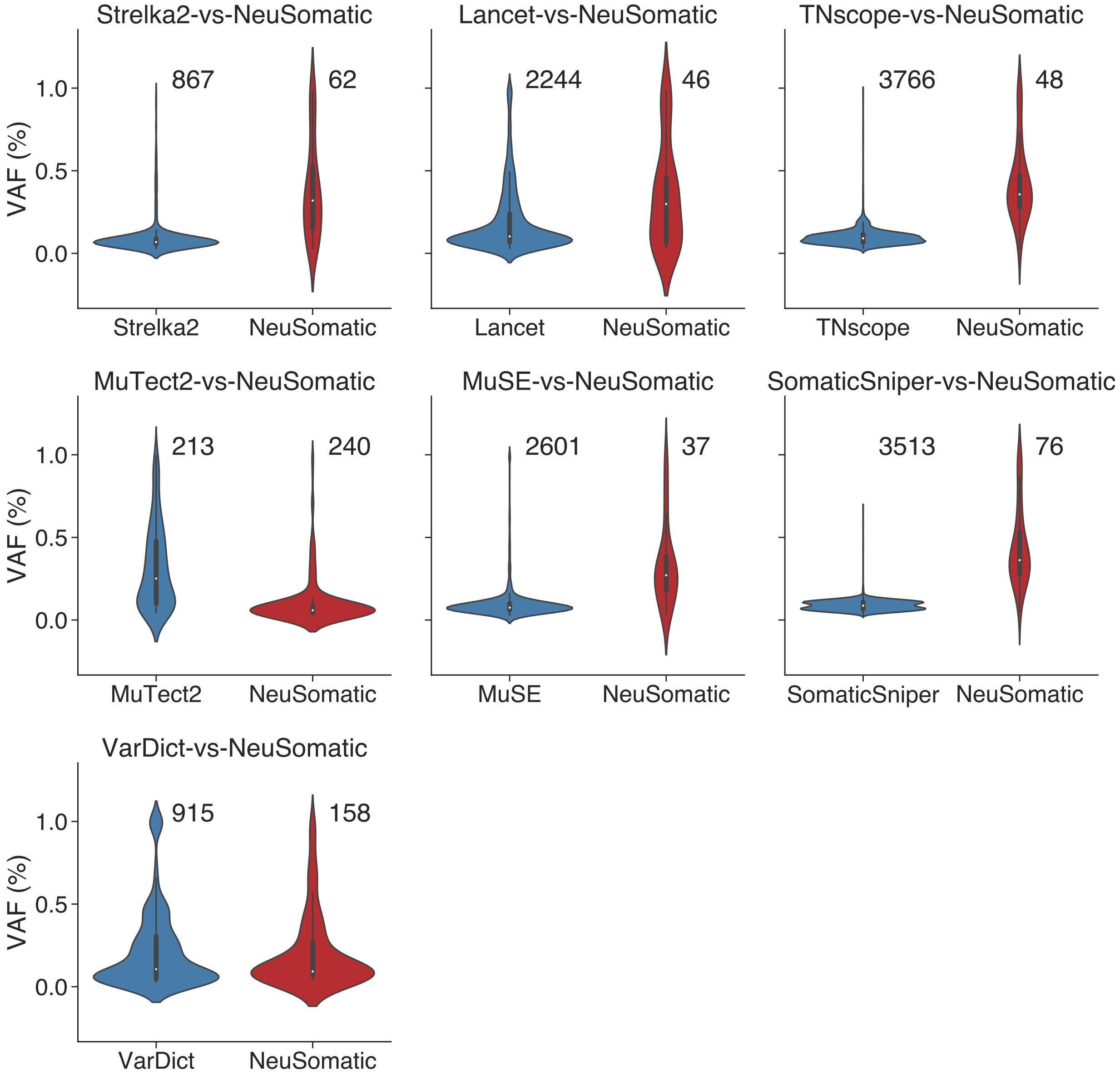


Supplementary Figure 14: The violin-plot comparison of the VAF distribution of private FN calls on WGS dataset. In each subfigure we compared one of the conventional somatic mutation detection schemes against NeuSomatic. For a $X$-vs-$Y$ subfigure, we identified $X_{\mathrm{FN}}$, the set of ground truth SNVs which were missed by algorithm $X$ (FN in $X$) in at least 11 out of 21 WGS replicates. Similarly, we identified $Y_{\mathrm{FN}}$, the set of ground truth SNVs which were missed by algorithm $Y$ (FN in $Y$) in at least 11 out of 21 WGS replicates. The figure then shows the VAF distribution of private FN calls for $X$ and $Y$. In other words, the violin-plot shows the VAF distribution of calls in the set $X_{\mathrm{FN}}/Y_{\mathrm{FN}}$ in blue and the VAF distribution of calls in the set $Y_{\mathrm{FN}}/X_{\mathrm{FN}}$in red. For most of the conventional schemes like Strelka2, the private FNs, which were correctly predicted by NeuSomatic, had low VAF which revealed their inferiority of such approaches in detecting low VAF mutations. In each $X$-vs-$Y$ subfigure, the number of private FNs for $X$ and $Y$ are reported on the top.
